# *PTEN* and *ARID1A* haploinsufficiency equip colonic epithelium for oncogenic transformation

**DOI:** 10.1101/2024.03.27.586939

**Authors:** Nefeli Skoufou-Papoutsaki, Sam Adler, Shenay Mehmed, Claire Tume, Cora Olpe, Edward Morrissey, Richard Kemp, Anne-Claire Girard, Elisa B Moutin, Chandra Sekhar Reddy Chilamakuri, Jodi L Miller, Cecilia Lindskog, Fabian Werle, Kate Marks, Francesca Perrone, Matthias Zilbauer, David S Tourigny, Douglas J Winton

## Abstract

Normal aged tissues are thought to exist as a patchwork of mutant clones. However, the relevance of driver mutations in normal tissue in terms of cancer initiation has not been well described. Here, we sought a quantitative understanding of how different cancer drivers achieve an age-related mutational footprint in the human colonic epithelium and to relate the clonal behaviours they generate to cancer risk. Metanalysis of contemporary multiregional sampling studies of colorectal tumours revealed many of the weak or moderate cancer drivers are trunk mutations present in the last common ancestor from which cancers arise. To study the processes by which such driver mutations could contribute to cancer predisposition, immunohistochemistry was used to detect PTEN, SMAD4 and ARID1A deficient clones in normal colon FFPE surgical resection samples (N=182 patients). Age-related changes in clone size and frequency identified positive biases in clone dynamics that acted to increase the mutational footprint for ARID1A and PTEN but not SMAD4. *In vitro* engineered monoallelic loss of *PTEN* and *ARID1A* implicated specific altered downstream pathways and acquired pro-oncogenic cellular fates corresponding to haploinsufficiency for these genes. *In situ* analysis confirmed enhanced proliferation in both PTEN and ARID1A deficient clones and creation of an immune exclusive microenvironment associated with ARID1A deficiency. The behaviours resulting from haploinsufficiency of *PTEN* and *ARID1A* exemplify how priming of the tissue through somatic mosaicism could contribute alternative combinations of genetic events leading to transformation.

## Introduction

The classical view of colorectal cancer (CRC) progression invokes a series of mutational events, starting with homozygous *APC* loss, mutations in *KRAS* and subsequently *TP53* (Fearon and Vogelstein, 1990). Late-stage events driving progression are recognised as subclonal as they occur within an established tumour mass (Gerstung *et al*., 2020). However, recent multi-regional sampling studies of CRCs have identified supposed late events in the trunk and early events in the branches of phylogenetic trees (Sottoriva *et al*., 2015; Uchi *et al*., 2016; Martincorena *et al*., 2017; Roerink *et al*., 2018; Saito *et al*., 2018; Banerjee *et al*., 2021; Househam *et al*., 2022). An interpretation of this finding is that the order of mutations is not prescribed and instead cancers arise due to reaching a “full-house” threshold of driver mutations within a single cell (Tomasetti *et al*., 2015; Williams *et al*., 2016).

It is recently appreciated that normal aged human tissues, including the colonic epithelium, are composed of a patchwork of cancer-driver mutations (Martincorena *et al*., 2015, 2018; Lee-Six *et al*., 2019). Such studies document the presence of these events with little quantitative insight into the mutational burden of specific cancer-drivers or the nature of the selection biases that create them. In addition, recent studies have demonstrated that pro-oncogenic clones in genetically engineered mouse models can have “supercompetitor” phenotypes altering the behaviours of neighbouring cells to promote their own selection (Flanagan *et al*., 2021; van Neerven *et al*., 2021; Yum *et al*., 2021). The behaviours of pro-oncogenic clones in normal human tissue have not been previously investigated. Consequently, the availability of driver clones in normal tissue and their capacity to contribute to cancer initiation remains unknown.

Here, we detect *in situ* clones in normal human colonic epithelium that are deficient for ARID1A, SMAD4 or PTEN, which are recurrently mutated tumour suppressor genes associated with the truncal events in CRC. Subsequent characterisation identifies clonal advantage and pro-oncogenic cellular behaviours associated with molecular and phenotypic haploinsufficiency for *ARID1A* and *PTEN* in normal tissue that facilitate and increase their availability for oncogenic transformation. This finding supports the idea that some tumour suppressor genes have haploinsufficient roles in carcinogenesis (Inoue and Fry, 2017) but extends their effect to a pre-neoplastic context, exemplifying how priming of the tissue through somatic mosaicism could contribute to cancer initiation.

## Results

### Metanalysis of truncal CRC drivers

Established driver mutations of CRC that are identified from extensive bulk sequencing of advanced cancers can be either clonal or subclonal events. To recognise the former as potential drivers involved in cancer initiation, a metanalysis of multi-regional sampling studies was performed (N= 115 tumours) (Sottoriva *et al*., 2015; Uchi *et al*., 2016; Roerink *et al*., 2018; Saito *et al*., 2018; Banerjee *et al*., 2021; Househam *et al*., 2022). Truncal driver mutations were identified based on the ratio of non-synonymous to synonymous substitutions (dN/dS >1) for CRC (Martincorena *et al*., 2017) (Fig. 1a, Supplementary Fig. 1) (Suppl. Data Table 1). The majority of tumours (70%) contained 2-4 truncal hits in recognised cancer driver genes irrespective of their *APC* status (Fig. 1b-c). While most of the single hit tumours were *APC*-driven, this accounted for only 3% of all tumours (Fig. 1d-e). Importantly, analysis of tumours with at least three driver mutations revealed that there is an equal number of those containing the classical CRC combination of *APC*, *KRAS* and *TP53* only and those that contained any other combination of mutations (Fig. 1f). More specifically, events that have been previously classified as late or subclonal from bulk studies, such as *SMAD4* or *PTEN* (Gerstung *et al*., 2020), were also found in the trunk of phylogenetic trees (Fig. 1g), either co-occurring with early events like *APC* or *KRAS*, or not. Collectively, these results highlight that weaker drivers are commonly implicated in early stages of cancer development.

**Figure 1:**
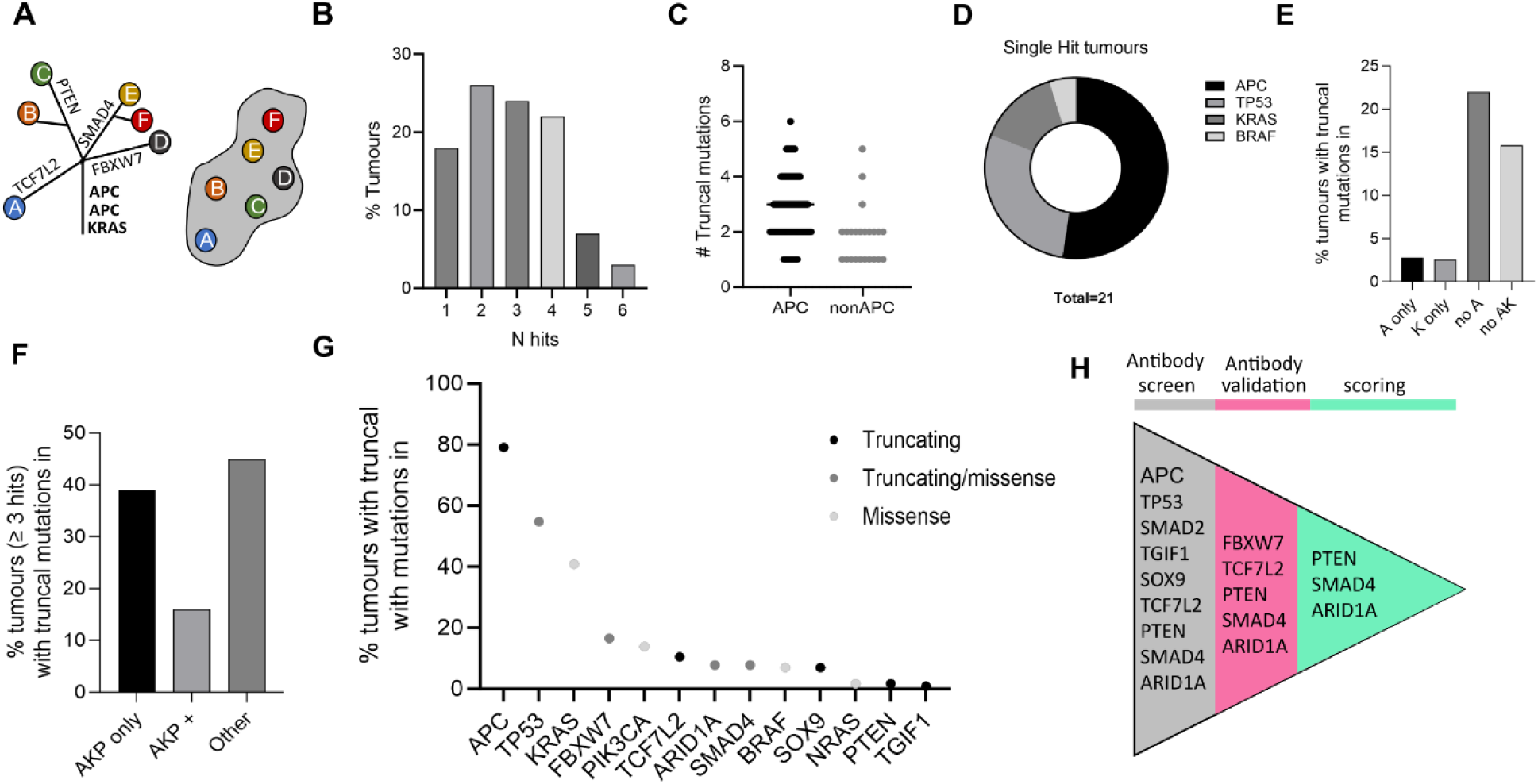
Trunk mutations from a metanalysis of multi-regional sampling studies. **(A)** Example of truncal analysis schematic. Grey mass represents tumour which has been sampled in 6 regions A-F. Mutations found in all parts of the tumour sampled, are placed on the trunk of the tree representing the last common ancestor. Other mutations define subclones within the tumour. **(B)** Distribution of number of truncal mutations derived from colorectal cancers (CRCs) for genes with ratios of dN/dS >1. **(C)** Number of truncal mutations in tumours with or without APC mutations. **(D)** Composition of single hit tumours. **(E)** Percentage of tumours driven by early selected events. A: APC, K: KRAS, AK: APC KRAS. **(F)** Combinations of mutations found in the trunk including tumours with three or more hits. AKP only: tumour with three hits in one or more of APC, KRAS or TP53; AKP+: tumours with four or more hits including one or more of APC KRAS or TP53 plus other driver(s), Other: any other combination of genetic events. **(G)** Percentage frequency of tumours with truncal mutations. Allows for identification of genes with recurrent truncating mutations. 8-G N=115 tumours. **(H)** Pipeline for identification of cancer driver genes for which loss of staining can be detected using immunohistochemistry. List in grey: candidates for antibody screen from (G) with truncating mutations. Pink indicates targets that were included in antibody validation screen. Antibodies for PTEN, ARID1A and SMAD4 in green passed all validation steps.

### Loss of staining in tumour suppressor genes *PTEN*, *SMAD4* and *ARID1A* detected in the human colon

Previously we used *in situ* immunodetection of X-linked genes in normal FFPE colonic tissue to benchmark neutral, and quantify biased clone dynamics to show that they dictate the mutational burden of specific events in the tissue (Nicholson *et al*., 2018; Olpe *et al*., 2021). Starting with the truncal drivers that were identified from the metanalysis, tumour suppressor genes in which encoded proteins are depleted by truncating mutations were screened with a panel of antibodies to determine the feasibility of adapting the approach to autosomal cancer-associated genes (Fig. 1g-h). Many antibodies failed the screen due to heterogenous or low baseline immunoreactivity within the epithelium precluding detection of clonal loss (Supplementary Fig. 2a-c). Five antibodies giving homogenous staining were run against an initial panel of 10 FFPE blocks taken from patients 60-80 years old and sectioned *en face*. Where candidate clones with reduced immunoreactivity that was focally restricted to one or multiple contiguous crypts were identified, antibodies were validated with tissues containing genetic knockouts of the relevant gene, and clones themselves validated by using independent antibodies in serial section (Supplementary Fig. 2d-e). The analysis indicated the presence of epithelial clones with reduced immunoreactivity for PTEN, SMAD4 and ARID1A (Figure 2a-c).

**Figure 2:**
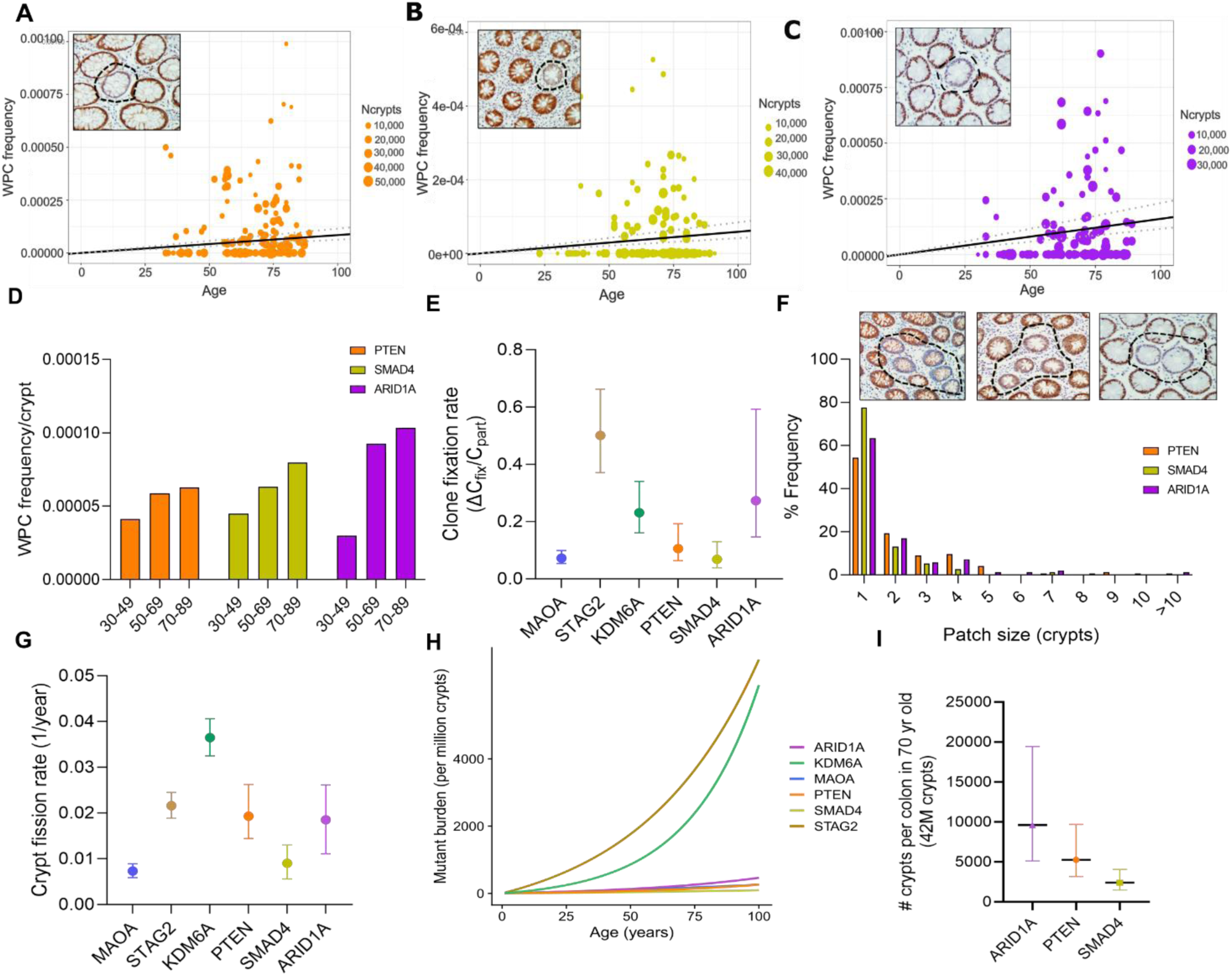
Clonal dynamics for tumour suppressor genes PTEN, SMAD4 and ARID1A. **(A-C)** Representative single Wholly Populated Crypts (WPC) images and plots of clone frequency against patient age for: (A) PTEN, Naa 103 patients, Naa 2,492,712 total crypts; (B) SMAD4, N “” 88 patients, Naa 1,661,059 crypts ; (C) ARID1A, Naa 101 patients, Naa 1,990,179 crypts. **(D)** Frequency of WPC across different age groups_ **(E)** Comparison of inferred intra-crypt dynamics (clone fixation rates) for genes shown. Calculated as ratio of slope of WPCs (ll.Cfix) versus frequency of Partially Populated Crypts (PPC) (Cpart) using mathematical modelling. **(F)** Patch size distribution and representative examples of PTEN, SMAD4 and ARID1A patches. **(G)** Comparison of inferred crypt fission (clonal expansion rates) per year for genes shown. Calculated based on the patch size distribution **(F)** using mathematical modelling_ **(H)** Mutational burden expressed per million crypts, incorporating biases in clone fixation and expansion rates. (I) Number of mutant crypts per colon in 70 year old individual. Error bars represent confidence intervals (Cl) 95%.

### ARID1A deficient clones exhibit advantage in clone fixation

To determine if these tumour suppressor deficient clones confer an advantage in clone dynamics, immunohistochemistry (IHC) was performed on FFPE sections from around 100 individuals (age range: 30-91) for PTEN, SMAD4 and ARID1A. Epithelial clones are permanently fixed within the colon when they successfully populate the whole crypt in the well characterised phenomenon of monoclonal conversion (Winton, Blount and Ponder, 1988; Winton and Ponder, 1990; Cook, Williams and Thomas, 2000). The frequency of fixed clones, wholly populated crypts (WPC), was measured for each patient and plotted as a function of age (Fig. 2a-c) (Suppl. Data Table 2-4). Partially populated crypts (PPCs) representing unresolved clones and multicrypt patch sizes were also recorded. Linear regression revealed an age-related increase in the number of WPCs for all three marks, with a slope, ΔC_fix_, of 8.45 x 10^-7^ for PTEN (95% CI: 6.26 x 10^-7^-1.15 x 10^-6^), 5.87 x 10^-7^ for SMAD4 (95% CI: 4.14 x 10^-7^-8.42 x 10^-7^ and 1.60 x 10^-6^ for ARID1A (95% CI: 1.15 x 10^-6^-2.31 x 10^-6^), with more WPCs found in older individuals (Fig. 2d).

The ratio of ΔC_fix_ over the frequency of PPCs (C_part_) allows for a quantitative description of the process of monoclonal conversion and the comparison of different clonal events independent of their mutation rate (Nicholson *et al*., 2018). PTEN and SMAD4 exhibited similar clone fixation rates with previously benchmarked neutral loss of MAOA. In contrast, ARID1A deficient clones were shown to have a positive bias in clone fixation similar to that previously described for the X-linked gene KDM6A (Fig. 2e) (Olpe *et al*., 2021).

### PTEN and ARID1A deficient clones exhibit bias towards clonal expansion via crypt fission

Clonal expansions generating multicrypt patches arise from crypt fission, the rates of which can be inferred from the patch size distribution (Fig. 2f). The homeostatic rate of colonic crypt fission has been previously described at around 0.7% of crypts undergoing fission per year (Nicholson *et al*., 2018). This is similar to the rate estimated here for SMAD4 deficient clones (Fig. 2g). In contrast, PTEN and ARID1A deficiency confers a bias in clonal expansion, showing around a 3-fold increase in the rate of crypt fission (Fig. 2g).

The combination of both clone fixation and expansion processes dictates the mutational burden of individuals at a given age (Fig. 2h). For a 70-year old individual, if scaled for the entire human colon which contains around 42 million crypts, there would be around 10,000 ARID1A deficient clones (mean: 9618, CI: 5124-19446), 5,000 PTEN deficient clones (mean: 5250, CI: 3179-9702) and 2,500 SMAD4 deficient clones (mean: 2406, CI: 1491-4044) (Fig. 2i).

SMAD4 was not considered further due to the neutral behaviour of deficient clones in both clone fixation and fission.

### Deficient autosomal clones are the result of monoallelic loss

For tumour suppressor genes, it is typically thought that the loss of both alleles is required for the development of a phenotype. Initially, to assess whether the changes in clone dynamics are likely to associate with biallelic or monoallelic gene loss, multiplex immunofluorescence (IF) was used to quantify the reduction in immunostaining intensity of deficient clones assigned to autosomal (*PTEN* or *ARID1A*) or X-linked (*MAOA*, *STAG2* or *KDM6A*) genes versus their adjacent WT crypt neighbours (Fig. 3a-b, Supplementary Fig. 3). As expected, due to functional hemizygosity of X-linked genes MAOA, STAG2 or KDM6A deficient clones showed an 80-90% reduction in immunoreactivity. In contrast, a 40-50% reduction in autosomal gene clones was observed, consistent with monoallelic loss.

**Figure 3:**
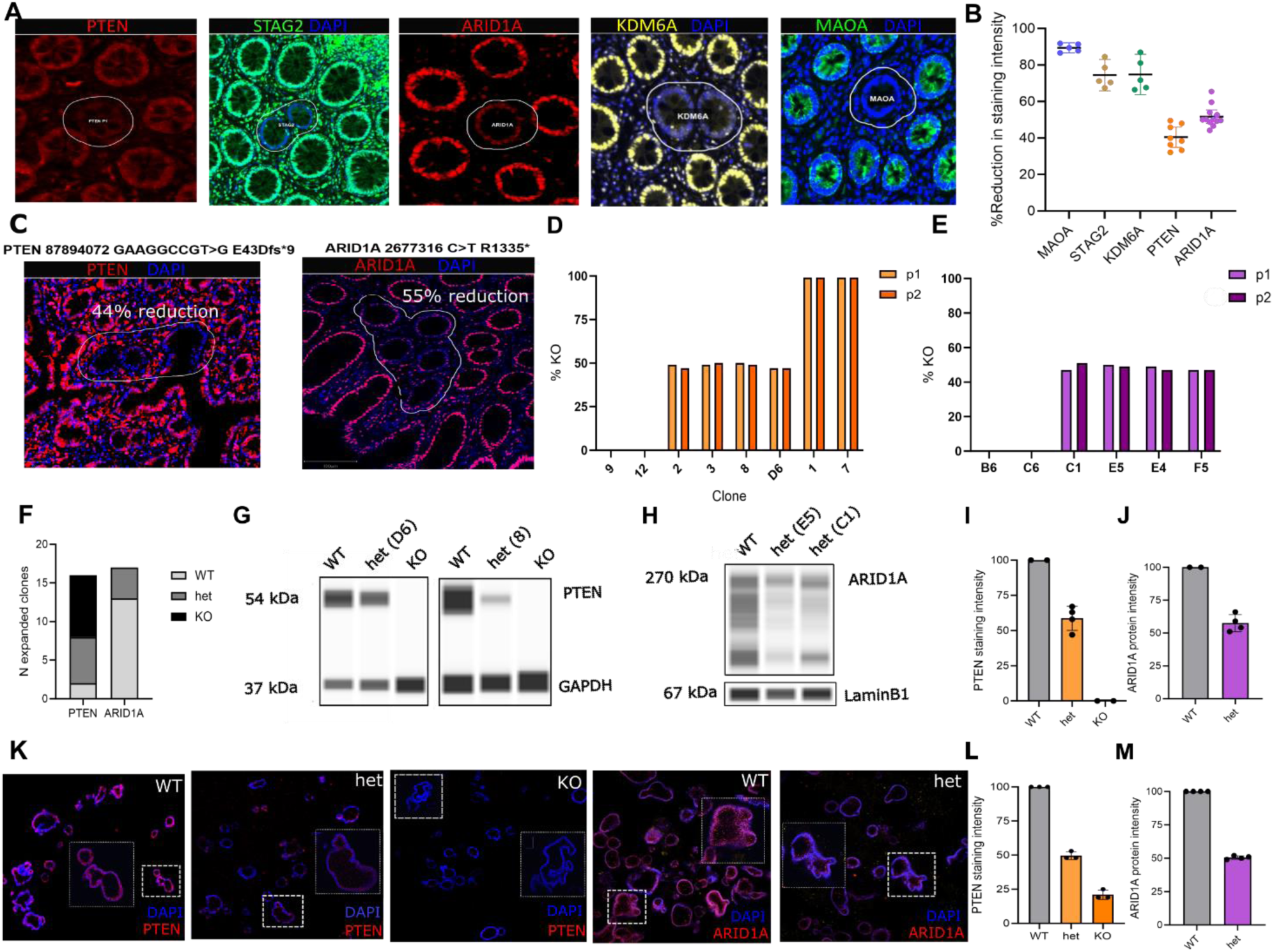
Quantification of protein expression of autosomal clones indicates monoallelic inactivation. **(A)** Representative examples of detected clones with immunofluorescence. **(B)** Quantification of staining intensity for clones deficient in proteins shown. **(C)** Left-Example of PTEN deficient patch with 44% reduction in staining intensity and a heterozygous PTE.lf frameshift mutation (E43Dfs*9). Right-Example of ARIDlA deficient patch with 55% reduction in staining intensity and a heterozygous ARIDlA truncating mutation (R1335*). **(D, E)** Percentage KO score for PTEN (D) and ARIDlA (E) for clones generated from single organoids. **{F)** Genotypes of generated clones and their proportions. **(G, H)** Wes Biotechne protein analysis for PTEN (G) and ARIDlA (H) clones. (I, J) Quantification of PTEN (I) and ARIDlA (J) staining intensity using Wes. **(K)** Wholemount organoid images stained for PTEN and ARIDlA and genotypes shown. **(L, M)** Quantification of signal intensity from imaging of organoid wholemounts stained for PTEN (L) and ARIDlA (M).

It has been previously suggested that while the transcriptome reflects copy number alterations, the proteome does not, tending to revert to a diploid state (Stingele *et al*., 2012). However, the link between gene dosage and protein abundance remains unclear. To more directly link PTEN and ARID1A deficient clones with monoallelic loss, a laser-capture microdissection and targeted amplicon sequencing strategy was employed on deficient patches. From this analysis, samples with identified truncating mutations in the coding region of *PTEN* and *ARID1A* correlated with a 50% reduction in protein intensity (Fig. 3c).

To confirm the effect of monoallelic loss on protein expression, CRISPR-Cas9 was used to introduce heterozygous mutations in PTEN and ARID1A in human intestinal organoids using a modified ribonucleoprotein (RNP) approach (Skoufou-Papoutsaki *et al*., 2023). Dead Cas9 protein was included in the RNP complex to limit the efficiency of genomic edits and generate heterozygous clones (50% KO) (Supplementary Fig. 4 a-c). These were isolated by picking individual organoids and identified by sequencing after expansion *in vitro*. The generated heterozygous clones contained one copy of a WT allele and one copy of an appropriately edited allele (+1, -7, -4) in an early exon. Isolation of isogenic controls of homozygous KO (100% KO) and WT clones (0% KO) was also attempted (Fig. 3d-e, Supplementary Fig. 4d, 4e). Although the editing efficiency was similar for *PTEN* and *ARID1A* (40% KO on population level), the WT:het:KO genotype ratio of the obtained clones was 2:6:8 and 13:4:0, respectively (Fig. 3f). The differences in frequency of clonal genotypes likely reflect their relative growth pattern, with *ARID1A* KO mutations incapable of clonal growth in an organoid context. Indeed, others have also shown that crypts isolated from *ARID1A* KO mice were unable to form spheroids (Hiramatsu *et al*., 2019). The observed genotype ratios might suggest that while homozygous KO might be the most advantageous PTEN genotype, organoid growth is more favourable in an ARID1A WT context.

Expanded *PTEN* and *ARID1A* heterozygous organoid clones were confirmed to have a 40-60% reduction in protein expression in comparison to WT clones as assessed by automated capillary immunoassay platform, Wes (Biotechne). In contrast, no staining was observed in PTEN homozygous KO organoids (Figure 3g-j). Mimicking the *in situ* detection used in patient samples, organoids were fixed and stained as a wholemount using immunofluorescence. This approach confirmed reduced staining in heterozygous *PTEN* and *ARID1A* mutant organoids (Fig. 3k-m).

Collectively, these findings demonstrate that monoallelic gene loss correlates with a 50% reduction at the protein level and that clones deficient in PTEN and ARID1A in the human colon arise due to monoallelic loss.

### *PTEN* and *ARID1A* heterozygous organoids acquire changes in downstream pathways

To explore how haploinsufficiency of *PTEN* and *ARID1A* is achieved at a molecular level, heterozygous organoids were further investigated. For PTEN, the impact of heterozygous mutations was assessed on the known downstream target p-Akt which was previously shown to be elevated with homozygous loss (Skoufou-Papoutsaki *et al*., 2023). A similar increase in the protein level of p-AKT was observed in PTEN heterozygotes (Fig. 4a). Interestingly, this increase inversely correlated with the level of PTEN protein expression detected in the generated clones.

**Figure 4:**
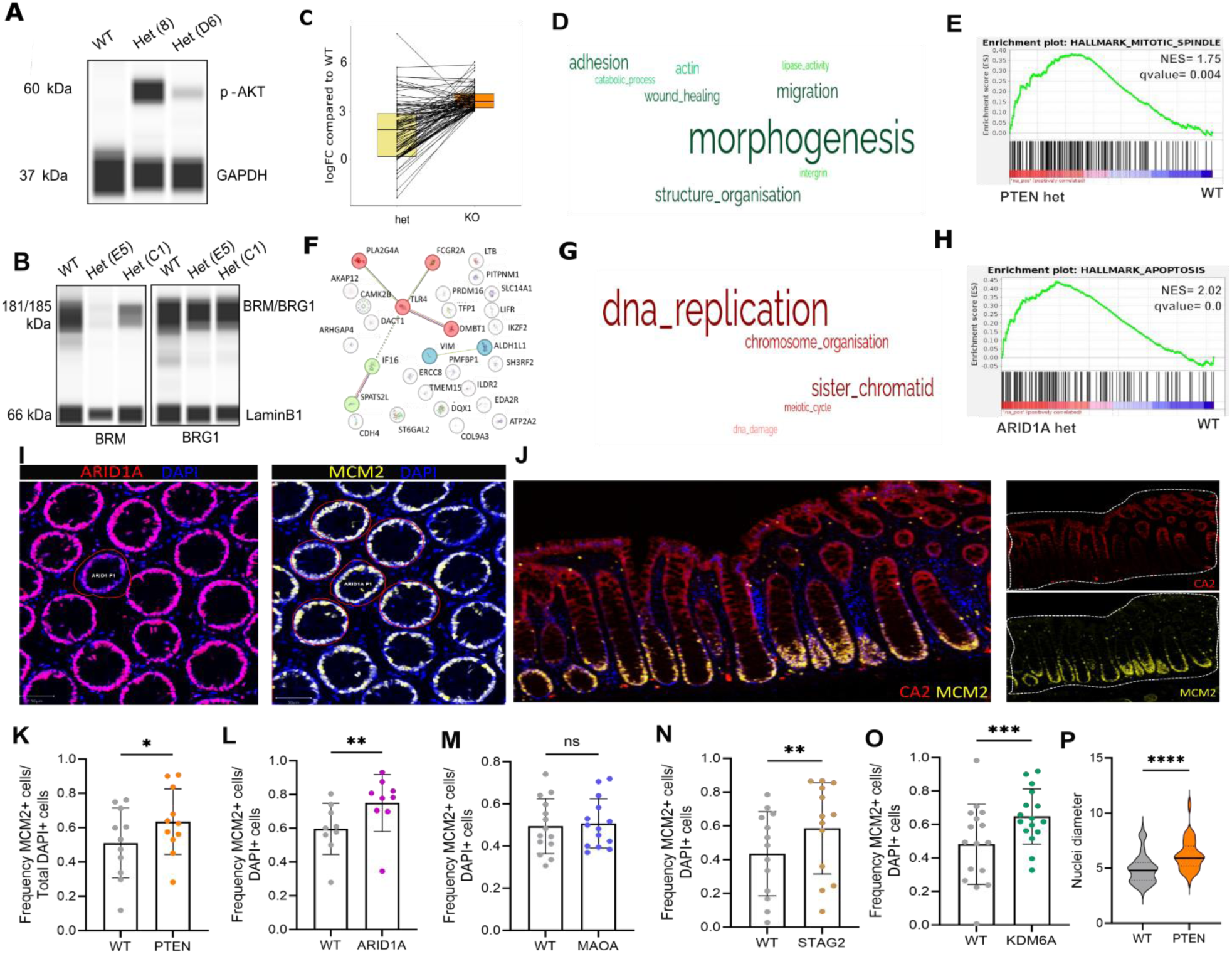
Characterisation of PTEN and ARID1A deficient organoids and crypt epithelium. **(A)** p-AKT Wes Biotechne on PTEN edited organoids. **(B)** BRM and BRGl Wes Biotechne on ARIDlA edited organoids. **(C)** Log Fold Change of het vs WT or KO vs WT for top 100 KO genes based on logFC. **(D)** Word cloud of Gene Ontology Biological Processes (GOBP) keywords from positively enriched gene sets in PTEN het vs WT organoids. Size of word indicates number of related gene sets in cluster. **(E)** Example of relevant hallmark pathway identified using Gene Set Enrichment Analysis (GSEA). Shows positive enrichment of PTEN heterozygotes vs WT. GSEA was performed for all genes detected in PTEN heterozygotes vs WT comparison. Genes were ranked based on fold change and standard error, showing gene sets with FDR q-value < 1%. **(F)** Interaction network of significant differentially expressed genes in ARIDlA heterozygotes vs WT using STRING. **(G)** Word cloud of GOBP keywords from negatively enriched gene sets in ARIDlA heterozygotes vs WT comparison. Size of word indicates number of related gene sets in cluster. **(H)** Example of relevant hallmark pathway identified using GSEA. Shows positive enrichment of ARIDlA heterozygotes vs WT. GSEA was performed for all genes detected in ARIDlA heterozygotes vs WT comparison. Genes were ranked based on fold change and standard error, showing gene sets with FDR q-value <1%. **(I)** Images show example of MCM2 scoring strategy. ARIDlA and MCM2 staining on serial sections allows MCM2 counts to be performed within the ARIDlA deficient clone and in five WT neighbouring crypts. **(J)** Markers used for the definition of crypt axis. CA2 expressed at crypt top and MCM2 expressed at crypt bottom. CA2+ MCM2-crypt top, CA2+ MCM2+ crypt middle, CA2-MCM2+ crypt bottom. **(K-0)** Quantification of MCM2+ cells in deficient clones or five WT neighbours within the crypt bottom. Expressed as frequency out of total DAPI+ cells in the crypt. (K) PTEN, (L) ARIDlA, (M) MAOA, (N) STAG2, (0) KDM6A. **(P)** Nuclei diameter size (µm) quantified in WT or PTEN deficient cells in FFPE tissue. Wilcoxon paired t-test was used in K-O. (K) p=0.0114, (L) p= 0.0039, (M) p= 0.455, (N) p=0.0018, (0) p= 0.0009. Mann-Whitney test was used in P. (P) p < 0.0001. * p < 0.05, ** p < 0.01, *** p < 0.001.

ARID1A is part of the SWI/SNF complex that acts to regulate the activity of catalytic ATPases BRM or BRG1 to drive chromatin remodelling. In ARID1A heterozygous organoids, the status of both effectors was assessed and a decrease in BRM but not BRG1 observed (Fig. 4b). The same finding has also been previously observed in a mixed population of ARID1A KO organoids (90% KO without clone selection) (Skoufou-Papoutsaki *et al*., 2023).

Bulk RNA-seq was performed to gauge the extent of altered gene expression associated with heterozygosity for PTEN and ARID1A (Suppl. Data Table 5). For PTEN, only 4 genes were found to be significantly differentially expressed (DEG) after multiple testing corrections (*PLCXD2*, *LXOL4*, *TNNC1* and lncRNA transcript ENSG00000260401, p < 0.05). Despite the small number of DEGs, the direction of the change observed in PTEN heterozygous transcripts seemed to follow that of homozygous KOs with a lower fold change in the former (Fig. 4c). Specifically, around 65% of the commonly detected genes (10217 out of 16090) displayed the same direction of change in gene expression. Consequently, gene set enrichment analysis (GSEA) was performed on all detected genes ranked by on their fold change and standard error, resulting in several pathways showing enrichment (FDR < 1%). Redundant gene ontology biological processes (GOBP) were removed using Revigo (Supek *et al*., 2011) and those remaining were clustered based on their relatedness (Supplementary Fig. 5a-b). This revealed that *PTEN* monoallelic loss resulted in a transcriptomic shift associated with developmental and regenerative processes, such as organ morphogenesis (Fig. 4d). Hallmark pathway GSEA suggested that the competitive advantage of *PTEN* heterozygous cells could be linked to increased proliferation as indicated by upregulation of the mitotic spindle pathway and metabolic changes like activation of fatty acid metabolism (Fig. 4e, Supplementary Fig. 5c-d). Enrichment of similar pathways has been described with homozygous loss of PTEN in organoids (Skoufou-Papoutsaki *et al*., 2023). Of note PI3K and mTORC1 activation has been previously reported in *PTEN* heterozygous endometrial organoids (Geurts *et al*., 2023).

Transcriptional profiling of heterozygous *ARID1A* organoids relative to wildtype identified 39 DEGs (p < 0.05) (Suppl. Data Table 6). Of these, several members of immune-response related clusters identified using STRING were strongly downregulated (log_2_FC -2 to -7) and included *IF16*, *TLR4*, *FCGR2A* or *PLA2G4A* (Fig. 4f). Similar pathway analysis of all the detected genes was performed for ARID1A het loss which was associated with increased apoptosis and a general downregulation of DNA replication processes (FDR < 1%) (Fig. 4g-h, Supplementary Fig. 6).

### PTEN and ARID1A deficient clones exhibit increased cell proliferation in primary tissue

The organoid and the FFPE tissue data revealed opposing phenotypes in response to ARID1A deficiency, with a selective advantage in the tissue but compromised growth *in vitro,* accompanied by transcriptomic analysis revealing a trend towards increased apoptosis and reduced DNA replication. To try and relate the molecular and pathway adaptations observed in organoids to the positive bias in clone fates *in vivo* both PTEN and ARID1A deficient clones were profiled for cellular changes using multiplex IF in FFPE sections. The proportion of cycling cells occupying the proliferative lower crypt region was determined based on expression of the DNA replication licensing factor MCM2 (Fig. 4i), Supplementary Fig. 7). Crypt levels were defined based on MCM2 and CA2 expression, with crypt bottom indicated as MCM2+CA2-(Fig. 4j). An increase in cell proliferation was seen in both PTEN and ARID1A deficient clones compared to their WT neighbours as indicated by a higher frequency of MCM2 positive cells (Fig. 4k-o). The same phenotype was seen in clones arising by loss of genes with biased clonal behaviours such as STAG2 and KDM6A but not in neutral MAOA deficient clones. This analysis suggests that clonal biases associate with increased cell proliferation. Notably this included an increase in proliferation in ARID1A deficient clones despite the adverse consequences of its deletion in organoids culture. In confirmation, no increase in cell cycle arrest (pH3+) or apoptosis (cleaved PARP) was seen in ARID1A deficient clones *in vivo* suggesting that context influences the epithelial phenotype (Supplementary Fig. 7g,7v).

mTORC1 activation, which is a consequence of PTEN loss, can lead to larger cells (Fingar *et al*., 2002). For a functional assessment of the PTEN deficient clones *in vivo*, nuclei diameter was used as a proxy for cell size (Edens *et al*., 2013). Accordingly, PTEN deficient cells exhibited larger nuclei compared to WT cells in the primary FFPE tissue sections, suggesting an involvement of mTORC1 in their phenotype (Fig. 4p). The concordance between the organoid data and the observed behaviours of the PTEN deficient clones *in vivo* is indicative of a cell-intrinsic selective advantage upon monoallelic PTEN loss.

### ARID1A deficient patches create microenvironment of immune exclusion

Motivated by the downregulation of immune response genes, and seeking an explanation for the apparent discrepancy implied by the compromised behaviour of *ARID1A* heterozygous knockout organoids and the positive biases and increased proliferation of ARID1A deficient clones in whole tissue, we sought to explore cross-talk with the surrounding stroma. Stromal cell profiling using multiplex IF was performed on ARID1A deficient clones using panels that visualised mesenchymal cells, T-cells, B-cells, dendritic cells, neutrophils and macrophages (Fig. 5a-c, Figure S8-9). ARID1A deficient patches were seen to exhibit an overall reduced number of immune cells (CD45+) in both the crypts and the stroma (Fig. 5d-e). More specifically, in terms of stomal immune composition there was a significant reduction of dendritic cells (CD11c+), B-cells (CD20+) and macrophages (CD68+) seen within the stroma of ARID1A deficient patches compared to matched WT patches of the same area (Fig. 5f-g, Supplementary Fig. 8g). Both types of TIM3+ cells, including T-cells (CD3+TIM3+), which can mark IFNγ secreting or exhausted T-cells (Das, Zhu and Kuchroo, 2017), and macrophages (CD68+TIM3+), in addition to CD163-macrophages (CD68+CD163-) showed a reduction within ARID1A deficient patches (Fig.5h-m).

**Figure 5:**
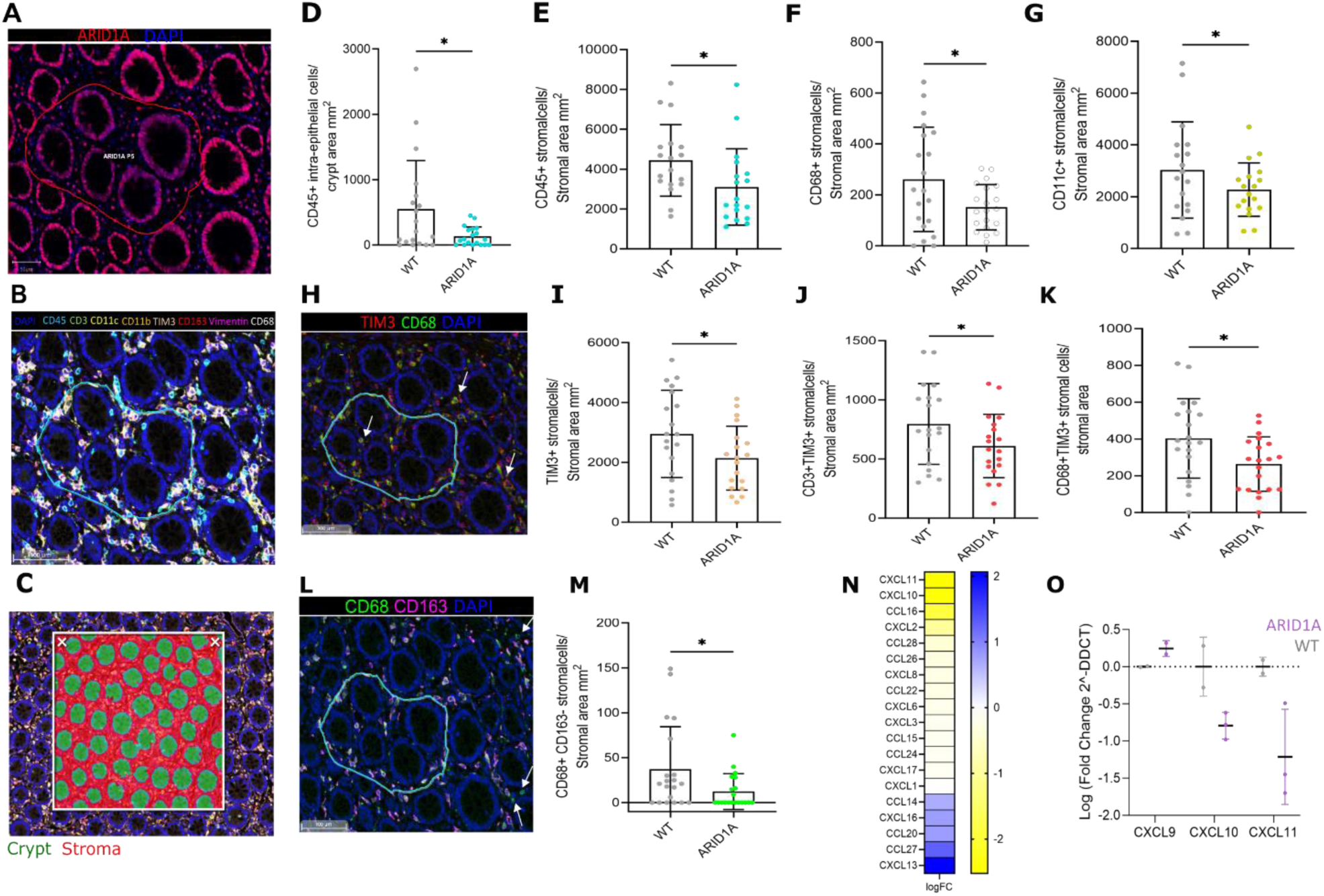
ARID1A deficient patches create an immune exclusion microenvironment. **(A)** Immunofluorescent image defining the area occupied by a patch of 5 deficient ARIDlA crypts and their associated stroma for downstream characterisation. **(B)** Staining of deficient ARIDlA patches for immune and mesenchymal cells using the markers shown. **(C)** Image of crypt and stromal areas assigned using random forest classifier training. **{D)** Quantification of CD45+ cells in crypt epithelium of ARIDlA patches (>5 crypts) and matched WT patch of same area. Normalised against crypt area. **(E-G)** Quantification of positive cells in the stroma associated with ARIDlA patches (>5 crypts) and matched WT patch of same area. Normalised against stromal area: (E) CD45 stroma; (F) CD68+ stroma; (G) CDllc+ stroma. **(H)** Co-localisation of CD68 and TIM3. TIM3+CD68+ cells indicated by white arrows. **(1-K)** Quantification of positive cells in the stroma associated with ARIDlA patches and matched WT patch of same area. Normalised against stromal area: (I) TIM3+ stroma ; (J) CD3+TIM3+ stroma; (K) CD68+TIM3+. **(L)** Co-localisation of CD68 and CD163. CD68+CD163-indicated by white arrows. **{M)** Quantification of CD68+/CD163-cells in the stroma associated with ARIDlA patches (>5 crypts) and matched WT patch of same area. Normalised against stromal area. **(N)** Heatmap of logFC of ARID1A heterozygous vs WT organoids for all detected chemokines. **(0)** qPCR on IFNy target chemokines on ARID1A heterozygotes and WT organoids. Wilcoxon paired t-test was used D-G, I-K, M. (D) p= 0.026 (E) p= 0.012, (F) p=0.034, (G) p= 0.044, (I) p= 0.017, (J) p=0.032, (K) p=0.032, (M) p = 0.049, * p < 0.05.

Relatedly a recent elegant study showed that immune evasion in *ARID1A* mutant tumours is mediated by impaired chromatin accessibility of IFNγ responsive genes including Th1 chemokines CXCL9, CXCL10 and CXCL11 (Li *et al*., 2020). Reviewing the RNA-seq data obtained for ARID1A heterozygous organoids revealed that although CXCL11 and CXCL10 were not significantly downregulated after multiple testing correction, they were the most highly downregulated detected chemokines (Fig. 5n). This was then directly confirmed using qPCR, with ARID1A heterozygous organoids showing an almost 2-fold reduction of CXCL10 and CXCL11 RNA expression (Fig. 5o). Importantly, in the study by Li and colleagues, the impact of reduced cytokine expression following *ARID1A* mutation was directly demonstrated by blocking the binding to their cognate receptor CXCR3 using an antibody, an intervention that acted to promote growth of colon cancer cells, highlighting a pro-oncogenic role of *ARID1A* mutations.

Overall, these results suggest that epithelial ARID1A deficiency acts non-cell autonomously to modulate the local immune environment and promote clonal expansion.

### Tissue representation defines oncogenic passengers and cancer-driver potency

To relate the increased tissue footprint arising from biased clonal behaviours to the incidence in cancers, the relative frequencies of genetic events in cancer and normal tissue were compared. The objective was to elucidate whether these genes potentiate cancer initiation, or are instead represented in CRC at a frequency that might be expected from the normal colon. As a motivating example, the mutational burden of the cancer driver *TP53* in normal oesophagus falls far below that observed in oesophageal cancer, while the mutational burden of *NOTCH1* is much higher in normal tissue, representing an extreme example of cancer-protective behaviour (Martincorena *et al*., 2018; Abby *et al*., 2023).

The comparison was performed by simulating mutational burden in normal colon by a given age using parameters (clone fixation and crypt fission rates) inferred for STAG2, KDM6A, PTEN, SMAD4 and ARID1A, and using publicly available data from COSMIC (Tate *et al*., 2019) to calculate the observed mutational burden for each of these genes in CRC (Fig. 6) (Suppl. Data Table 7-8). MAOA was not included because of its low event incidence in CRC. STAG2 was the only gene for which the predicted mutational burden in normal tissue reached that observed in CRC within an individual’s lifetime (69 years; range 13-110 years), suggesting that the frequency of *STAG2* events in CRC can be almost entirely explained by a positive selection bias in normal tissue, classifying it as an oncogenic passenger. Removing either the bias in clone fixation or crypt fission reduced the predicted lifetime mutational burden for *STAG2* to a level below that observed in CRC (Fig. 6a), which mimics the behaviour expected for a weak driver event, and similar to that seen with *KDM6A* (Fig. 6b).

**Figure 6:**
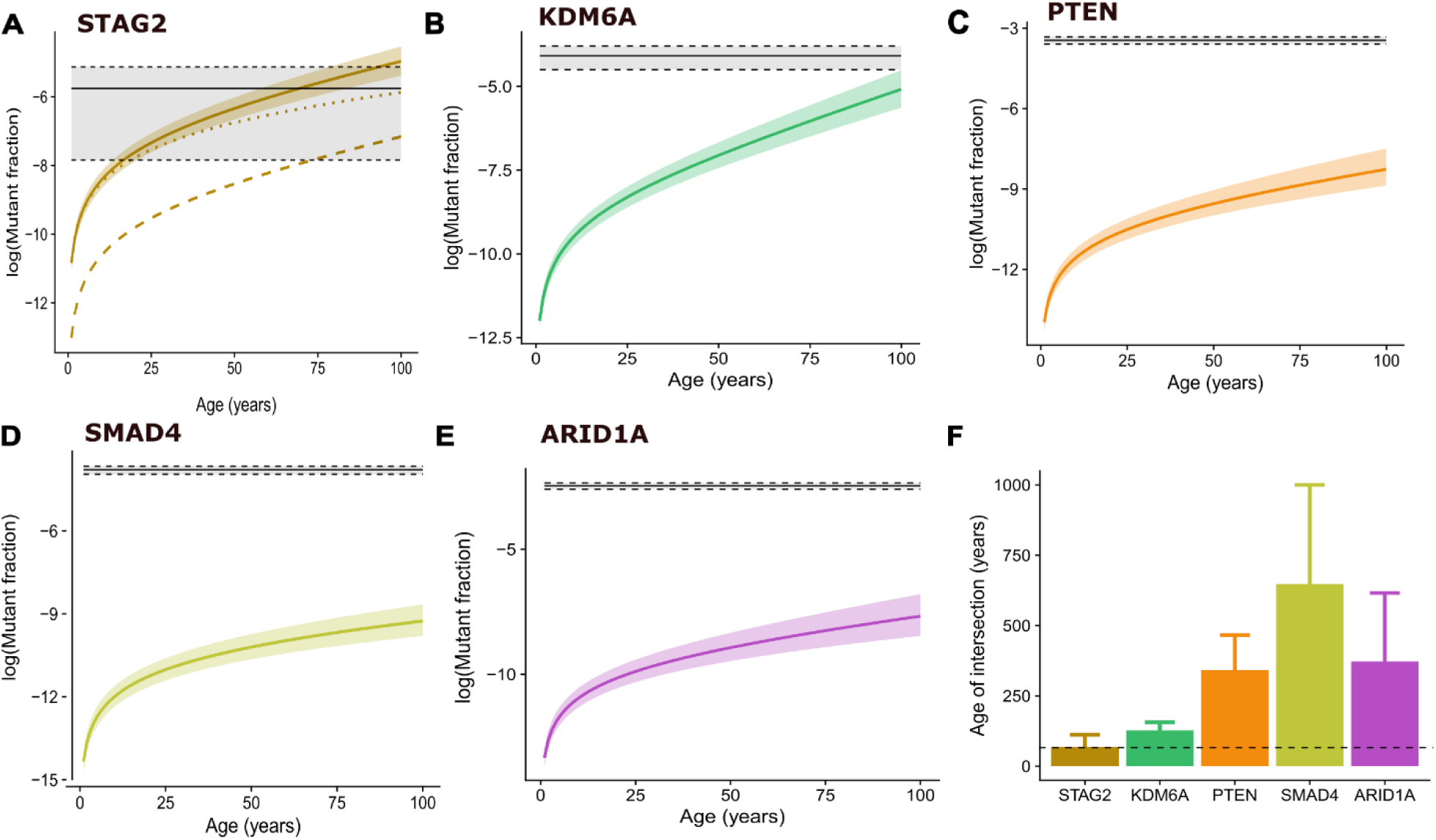
Comparison of tissue and cancer representation defines oncogenic passengers and drivers. **(A-E)** Predicted mutational burden in normal tissue based on clone dynamics (clone fixation and crypt fission rates) compared to frequency of truncating mutations for that gene in colorectal cancer in COSMIC. Shading indicates 95% Confidence Intervals (CI). Solid coloured line indicates mutational burden with fixation and fission bias. (A) STAG2. Upper dotted line indicates STAG2 burden without fission bias and lower dotted line indicates STAG2 burden without fixation bias ; (B) KDM6A ; (C) PTEN ; (D) SMAD4; (E) ARIDlA. **(F)** Predicted age of intersection of tissue burden and cancer burden. Dotted line indicates 70 years (within lifetime). Error bars indicate 95% CI.

In contrast, the predicted mutational burdens in normal colon over a lifetime fell far below that observed in CRC for *ARID1A*, *PTEN* and SMAD4 (Fig. 6c-e). Overrepresentation of these events in CRC therefore cannot be explained by the behaviour of the corresponding clones in normal tissue, since it is predicted to take within the range of 300-700 years for the normal mutational burdens to reach levels comparable to those observed in cancer (Fig. 6f). It is also seen that because of the neutrality of SMAD4 clones it takes many fold longer for the predicted mutational burden to reach that observed in cancer than it does for non-neutral ARID1A and PTEN, even though SMAD4 is represented less frequently in CRC that the other two genes (95% confidence interval (CI) [1.9-2.6%] for SMAD4 compared to [7.4-9.5%] and [2.8-3.6%] for ARID1A and PTEN, respectively).

Studying the mutational footprint of cancer associated genes in the normal tissue can allow reclassification of passenger and driver events. Although the increased mutational burden is not solely a feature of driver clones, it could act as a mechanism to increase their availability in normal tissue and subsequent probability for transformation. This, in addition to altered cellular changes induced by driver clones, demonstrates how priming of aged tissues, through somatic mosaicism, could equip the tissue for cancer transformation.

## Discussion

Recent studies have demonstrated that cancer driver mutations are frequent events in histologically normal human colon (Martincorena *et al*., 2015, 2018; Suda *et al*., 2018; Brunner *et al*., 2019; Lee-Six *et al*., 2019). Metanalysis of contemporary multiregional analyses of colonic cancers demonstrated that many of the “long tail” of weak or moderate cancer drivers are trunk mutations present in the last common ancestor from which cancers arise. Modelling of mutation rates based on genome sequencing data suggest that as few as three sequential mutations (Tomasetti *et al*., 2015) are sufficient to explain the incidence rates of colorectal cancers. It is also established that positive bias in clonal expansion is not a behaviour solely conferred by cancer drivers but is shared by mutations in genes that are not associated with oncogenesis in their resident tissue (Martincorena *et al*., 2018). These observations indicate a high degree of somatic variegation for cancer driver events in ageing tissues that has yet to be fully described and is of uncertain relevance for cancer initiation. Here we sought to extend our quantitative understanding of how different cancer drivers achieve an age-related mutational footprint in the human colonic epithelium and to relate the clonal behaviours they generate to cancer risk.

Clones deficient in PTEN, SMAD4 and ARID1A were readily detected by immunohistochemistry. Mathematical modelling established the clone dynamics that dictate their mutational burden. *In vitro* and *in situ* profiling of PTEN and ARID1A deficient clones pinpointed specific altered downstream pathways and cellular states that arise due to monoallelic loss. Using clone dynamics to establish the changing age-related degree of representation in normal tissue versus cancer provides a quantitative expression of driver potency and a conceptual framework enabling drivers and passengers to be classified. For example, the degree of representation in normal tissue of STAG2 deficient crypts that arises from positive bias in clonal fates, compared to that observed in cancer, revealed that the *STAG2* frequency in CRC can be solely explained by the increased mutational burden in normal tissue. Of note, using targeted sequencing Lee-Six *et al*. (2019) detected a proportion of crypts containing *STAG2* truncating mutations in normal colon from 50-60-year-old individuals that aligns well with the mutational burden estimated here by age 55 (95% CI [-0.05%, 0.34%] from targeted sequencing versus [0.14%, 0.29%] from simulation). Here, we demonstrate how pro-oncogenic events alter normal tissue maintenance processes to increase their mutational burden in the tissue and induce cellular changes that could predispose to cancer development.

While *STAG2* has a high mutational burden in normal tissue and lower in cancer, compared to the other tumour suppressor genes, *SMAD4* has the opposite phenotype of lower mutational burden in normal tissue but enrichment in CRC. Recent studies reconstructing the timing of CRC events assign SMAD4 mutation to a late stage of progression as it is predominantly found to be subclonal (Gerstung *et al*., 2020; Cornish *et al*., 2022). Despite this, the occasional presence of truncal *SMAD4* mutations might be indicative of a weak driver which relies on additional synergising events for transformation but does not employ a probabilistic strategy of increasing its tissue representation for co-occurrence.

Truncal *PTEN* mutations similarly suggest that *PTEN* mutations may also be implicated in early stages of cancer development. However, PTEN deficiency confers a biased expansion of clones that acts to increase the mutational footprint within the epithelium, with twice as many deficient crypts as SMAD4 by age 70. Haploinsufficiency in PTEN expression results in molecular activation of AKT signalling and a cellular proliferative phenotype that is potentially pro-oncogenic but that remains associated with a normal epithelium. It appears that monoallelic *PTEN* loss promotes a cancer-initiating potential by increasing its mutational footprint and increasing the probability of secondary synergising events on which it is dependent.

ARID1A deficient clones in addition to their ability to increase their mutational burden by positive biases in fixation and lateral expansion, also modulate the surrounding immune microenvironment towards immune exclusion. This phenotype of ARID1A mutations has been previously shown in ovarian and colorectal cancers and is mediated by downregulation of IFNγ responsive chemokines that include CXCL10 and CXCL11 (Li *et al*., 2020). Counterintuitively, this study established that this effect of ARID1A mutations is driving cancer immune evasion and promoting cancer growth. The creation of an immune privileged environment can be related to the role of *ARID1A* mutations as a driver of colitis-associated colorectal cancer and the selection of such mutations in the non-dysplastic epithelium of patients with ulcerative colitis (Kakiuchi *et al*., 2020; Olafsson *et al*., 2020). Given the relative difficulty in establishing and maintaining organoid cultures lacking ARID1A (Hiramatsu *et al*., 2019), it seems likely that the observed positive biases in clonal behaviours in the FFPE tissue is explained by increased immune fitness rather than enhanced proliferative fitness. The recognised cancer hallmark of “avoiding immune destruction” (Hanahan and Weinberg, 2011) is therefore simply and immediately acquired in the normal epithelium following the loss of function of a single ARID1A allele. Together, these observations suggest that the subsequent level of immune fitness conferred by this phenotype will scale with the severity of the local inflammatory environment.

The loss of one allele was sufficient for molecular or phenotypic changes in PTEN and ARID1A deficient clones. Although the haploinsufficiency of tumour suppressor genes is a well described phenomenon, especially in the context of cancer development (Inoue and Fry, 2017), we report phenotypic haploinsufficiency in normal human tissue. This phenotype can also extend to major drivers in CRC such as *APC* or *TP53* (Amos-Landgraf *et al*., 2012; Vermeulen *et al*., 2013; Preisler *et al*., 2021). In fact, the truncal analysis of multiregional sampled tumours revealed that 65% of all *APC* mutated tumours were driven by monoallelic loss. Estimates that three sequential mutations are sufficient to explain the incidence rates of colorectal cancers are agnostic to the nature of these events and deviation from stereotypical adenoma formation, due to biallelic inactivation of *APC* and monoallelic activation of *KRAS* (Fearon and Vogelstein, 1990), may be commonplace. Examples of such tumours are observed in the trunk of the tumour phylogenetic trees. More specifically, there was an equal number of tumours comprised of “strong” driver events (*APC*, *KRAS* and *TP53*) and tumours where “weaker” drivers were present, consistent with a degree of interchangeability in driver events. This would imply that instead of a prescribed combination of events there are “wild card” spots in carcinogenesis that are occupied based on the behaviour of clones in normal tissue. Our results position and contextualise *PTEN* or *ARID1A* mutations in the landscape of CRC initiation. Similar mechanisms could be employed by other weak drivers found in normal tissues to predispose to cancer development. An appreciation of how normal tissue maintenance processes can be subverted to increase the bioavailability of cancer drivers and how such drivers display cancer hallmark strategies from their inception is necessary to understand the clonal choreography that explains the incidence of colorectal cancer.

## Acknowledgements

We would like to thank the Histopathology, Genomics and Bioinformatics Core at the CRUK Cambridge Institute for their help with the project.

## Methods

### Human Tissue

Histologically normal colon tissue samples were obtained from cancer patients (Cancer Normal) from Addenbrooke’s Hospital Cambridge, Norwich University Hospital and St James University Hospital Leeds under full local research ethical committee approval (REC 15/WA/0131, 06/Q0108/307 & 08/H0304/85 as well as 12/LO/1217, respectively) according to UK Home Office regulations. Colectomy specimens were fixed in 10% neutral buffered formalin. From area s of tissue without macroscopically visible disease, mucosal sheets were removed from the specimens and embedded *en face* in paraffin (FFPE) blocks.

FFPE sections for staining were cut at 5 μm. To perform Laser-Capture Microdissection, sections were at 10 μm thickness onto special slides with PET membrane (Zeiss, 15511306) that had been irradiated with UV for 30 minutes prior to the cutting.

### Immunohistochemistry/Immunofluorescence Protocol

Immunohistochemistry was performed as previously described (Nicholson et al., 2018) (Suppl. Methods Table 1).

The immunohistochemistry protocol was adapted for immunofluorescence using secondary Alexa fluorophore conjugated antibody (1:200, Thermofisher) and DAPI (1 μg/mL). Slides were scanned on the Akoya Biosciences Vectra Polaris Automated Quantitative Pathology Imaging System (Perkin Elmer).

### Multiplexing immunofluorescence

Here, multiplexing immunofluorescence refers to the immunofluorescence approach where multiple cycles of staining and antibody stripping are performed allowing staining of antibodies of the same species. Three different multiplexing protocols were used: the 3-plex standard tyramide signal amplification (TSA), the 6-plex TSA with Opal dyes (Akoya) 6-plex and the 8-plex Signal Star (CST). For the standard TSA, the dewaxing and antigen retrieval were performed as previously described (Nicholson et al., 2018). All incubation steps were performed at room temperature. The primary antibody was incubated for 30 minutes. The secondary antibody was Horseradish Peroxidase (HRP) conjugated (Thermofisher anti-mouse T20916, anti-rabbit 65-6120 and anti-goat A15999, all 1:200) and was incubated for 30 minutes. TSA reagent conjugated to standard fluorophores (Cy5 Akoya (TS-000203), Cy3 Akoya (TS-000204) and Alexa Fluorophore 488 Thermofisher (B40953)) was added for 15 minutes (Table 4). That marked the completion of a staining cycle and antibody stripping was performed at 90 °C in a waterbath for 20 minutes. After the last TSA cycle DAPI (1 μg/mL in TBS) was added for 10 minutes followed by final washes with PBS for 10 minutes and 2 times and 15 minutes wash with TBS-T (Suppl. Methods Table 2).

For the TSA using Opal dyes, the Opal 6-colour manual detection kit (NEL811001KT) was followed as per the manufacturer’s instructions. The only modifications were that the first antigen retrieval was performed as previously described using a pressure cooker and that a waterbath set at 98 °C was used for each stripping cycle.

For the Signal Star, the SignalStar™ Multiplex Immunohistochemistry Assay (Cell Signalling Technologies) for Use on BOND RX Fully Automated Research Stainer (Leica Biosystems) was followed as per the manufacturer’s instructions.

### LCM of PTEN and ARID1A deficient patches

LCM was performed as described in Olpe et al., 2021 except from the modifications below. Fresh sections were stained using IHC for the mark of interest, PTEN or ARID1A, the slides were scanned on Leica Aperio AT2 and the location of the patch was mapped on the scanned slide. LCM was performed on serial section of the stained slide after dewaxing using Leica Multistainer ST5020. WT crypts were also captured to increase the input of DNA. The ratio of deficient crypts to total crypts captured was always 1:2.5 and the minimum patch size captured was total 10 crypts. Captured crypts were lysed in a volume of 12μl of Proteinase K buffer from the Arcturus PicoPure DNA kit (ThermoFisher, KIT0103).

### Fluidigm Assay Design

The gene of interest*, PTEN* or *ARID1A* was imported in the Fluidigm D3 Assay Design and dual coverage primers were designed. There were 45 amplicons used to cover *PTEN* and 114 to cover *ARID1A* and they were each pooled in 8 different multiplexing primer pools (Supplementary Methods Table 3-4).

### Library Prep using LP 8.8.6 Chip and Juno system

Due to the high genomic DNA to mastermix ratio in the LP 8.8.6 chip and the non-purified nature of the DNA lysed using the PicoPure method, an additional Ampure XP bead cleanup (Beckman Coulter™) was required prior to the first PCR. The bead to DNA ratio used was 1.8x to clean up the genomic DNA. Then, the library prep was performed using the LP 8.8.6 integrated fluidic circuit (IFC) (101-7663) according to the manufacturer’s instructions (Advanta NGS Library Preparation with Targeted DNA Seq Library Assays on the LP 8.8.6 IFC protocol). The only modification was the use of 17 cycles during the first PCR to accommodate for the low DNA input. Samples were submitted for an Illumina NovaSeq run for 150 bp paired end reads.

### Mutation calling in deficient patches

The SNV mutation calling was performed as mentioned below. Paired end assembler for Illumina sequences, PANDAseq was used to merge corresponding forward and reverse reads into an *in silico* amplicon (Masella *et al.,* 2012). The *in silico* recreated amplicons were then filtered to retain those that began and ended with the expected primer sequence and were the correct overall length. A Perl script was then used to count the number of reads for individual nucleotide (A, C, G or T) at each position in each amplicon. Samples with <500 reads were excluded from downstream analysis. A minimum of 1% VAF was set as the lower detection threshold. The mean variant allele frequency (VAF) and standard deviation was calculated for each position for all nucleotides in all amplicons. The single nucleotide variant (SNV) mutations were then called if the VAF of a particular sample was 3.29 standard deviations above the mean VAF in that specific position (p value < 0.001) for each amplicon independently. Only calls that were made in both replicate amplicons were considered. Only calls with two or more variant reads were included. In addition, the presence of insertion and deletions (indels) was also investigated using an alignment-based method, Amplicon-seq (https://github.com/crukci-bioinformatics/ampliconseq). The analysis was applied on mutation calls >2% VAF. Mutations that were called in three or more independent samples were excluded.

### Inference of clone dynamics

Clone and crypt counting using DeCryptICS was performed as previously described (Nicholson et al., 2018). The stem cell dynamics and fission rate associated were mathematically modelled as previously described (Nicholson et al., 2018) using the RHClones package (https://github.com/ElEd2/RHClones).

### Organoid culture

Intestinal mucosal biopsies were collected from sigmoid colon from children under 16 years old undergoing diagnostic colonoscopy at Addenbrookes Cambridge Hospital. This study was conducted with informed patient and/or carer consent as appropriate, and with full ethical approval (REC-12/EE/0265). Organoids were cultured as previously described (Skoufou-Papoutsaki et al., 2023). The only modification to the culture conditions was the addition of synthetic fusion Wnt (N001, IpA Therapeutics).

### CRISPR editing of organoids

A ribonucleoprotein approach was used for the CRISPR editing of human intestinal organoids as previously described (Skoufou-Papoutsaki et al., 2023) with the addition of an optimised ratio of dead (d) to active (a) Cas9 for each guide RNA. For PTEN the optimised ratio was 20:80 a:dCas9 while for the ARID1A guide the ratio was 50:50 a:dCas9. After electroporation, single organoids were picked using an IVF EZ-Grip pipette (Cooper Surgical, 7-72-2800 and 7-72-2155/1, 7-72-2200/1, 7-72-2290/1 for tips) and a dissecting microscope which were single cell dissociated and placed in individual wells allowing for clonal selection. Lines with pure 0, 50 or 100% editing as assessed by ICE Synthego (Conant *et al*., 2022) were expanded and subjected to downstream analysis.

### Automated Wes analysis

Protein extraction and automated Wes (Biotechne) analysis was performed as previously described (Skoufou-Papoutsaki et al., 2023).

### Organoid wholemounts

The organoid wholemount staining protocol was used as previously described (Skoufou-Papoutsaki et al., 2023).

### RNA extraction and bulk RNA-seq

RNA extraction and RNA-seq library prep was performed as previously described (Skoufou-Papoutsaki et al., 2023).

For the data analysis, Gene Set Enrichment Analysis (GSEA) was performed using Msigdbr on all the detected genes ranked based on their logfold change and standard error. Enriched pathways with an FDR <1% were considered. Revigo was used to identify redundancy in Gene Ontology Biological Processes (GOBP) gene sets and plot their relationship and interactions. Cluster representatives (terms remaining after redundancy reduction) were plotted in two-dimensional space by applying multidimensional scaling to a matrix of the GO term semantic similarities. Word clouds were created based on the Revigo scatter plot and simplification of the GO terms that most accurately describe the cluster in a biologically meaningful way. When necessary, synonym clusters that describe similar biological process were merged for simplicity (e.g. substrate adhesion and cell adhesion merged to create adhesion cluster). STRING (Szklarczyk *et al*., 2015) was used to identify interaction networks for the DEGs in ARID1A heterozygous vs WT organoids.

### Image analysis

Image Analysis was performed on Quantitative Pathology & Bioimage Analysis (QuPath) and Indica Labs Halo Image Analysis Platform for the FFPE tissue and Fiji for the organoid wholemounts. Fluorescence intensity of staining-deficient crypts and five WT neighbouring crypts was calculated on QuPath. For proliferation/differentiation markers, positive cells were manually counted in staining-deficient crypts and five WT neighbouring crypts in MCM2+CA2-areas (crypt bottom).

For the immune cell analysis on Halo, a random forest crypt-stroma classifier was trained and a cell segmentation algorithm (Module: Indica Labs HighPlex FL v4.1.3) was used to count the number of positive cells in either the crypt or stromal area. Only ARID1A patches of >5 deficient crypts were included in the analysis. For the Opal analysis, cells were counted within the patch or in a radius of 300 μm around the WT patch. For the Signal Star analysis, cells were counted within the patch and an adjacent (>300 μm away) randomly selected WT area of the same size. Counts were normalised for crypt or stromal area.

For the organoid wholemount analysis, the mean fluorescence intensity of each organoid was measured using the measure function in Fiji. The background signal and bleaching from the DAPI channel was subtracted by measuring the fluorescence intensity in areas with no signal or areas with DAPI only signal.

### Stem cell dynamics and crypt fission inference

Stem cell dynamics and fission rates associated with markers were mathematically modelled as previously described (Nicholson *et al*., 2018). A minor modification to the hierarchical model for fission rates was made by instead drawing patient-specific fission rates from a Student’s t-distribution according to

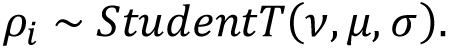

Here *ρ_i_* is the fission rate for patient *i* and the priors for population parameters ρ, σ and *v* are

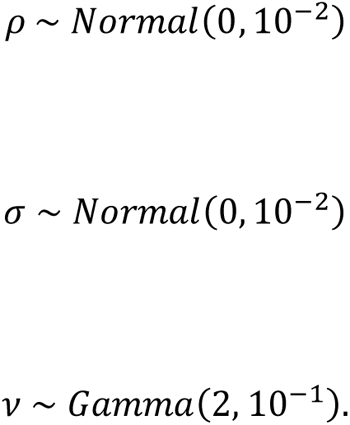

Using a Student’s t-distribution in place of a Normal distribution allows for control of heavier tails in the hierarchical model for patient-to-patient variability through the parameter *v*. Inferred values for *v* were spread across a wide range (0.31-148.12) suggesting that fission rate priors for different genes can display a substantial departure from the standard normal distribution, including regimes where the variance is not strictly well-defined.

### Estimated mutational burdens and age of intersections

Inferred stem cell dynamics and fission rates population parameters were used to calculate the estimated mutational burden in normal colon *B*(*t*) at a given age *t* according to the formula

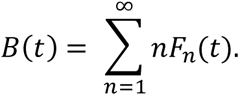

Here *F_n_(t)* is the distribution for the probability of a patch of mutated crypts of size *n* at age *t* calculated from population estimates for the slope of monoclonal accumulation Δ*C_monoclonal_* and fission rate ρ for that marker given by

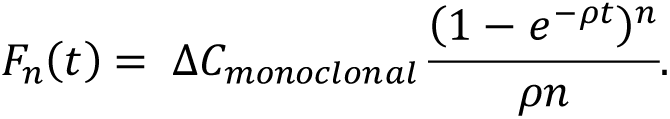

In practice, the infinite sum for *B*(*t*) was replaced by a finite sum over an extremely large number (in this case 500,000). 95% confidence intervals (CIs) for *B*(*t*) were calculated using the same formula replaced with 95% CIs on population parameter value estimates Δ*C_monoclonal_* and ρ for each marker.

To compare estimated mutational burden in normal epithelial with observed mutational burden in CRC, the proportion of colorectal tumours (primary samples not cultured) containing a truncating mutation (i.e. substitution-nonsense, frameshift, deletion-frameshift, insertion-frameshift, whole gene deletion) were extracted from the COSMIC database for each marker gene. The **age of intersection** was then defined at the age *t*^∗^ at which *B*(*t* ^∗^ ) equals the observed CRC mutational burden for that marker. Error bars displayed in the corresponding plots for age of intersection are then calculated from the intersections of the upper (respectively, lower) CI for *B*(*t*) with the lower (respectively, upper) 95% CI for the observed CRC mutational burden. Note that this representation of errors for estimated ages of intersection is overly stringent in the sense that comparing overlapping error bars is too conservative a criterion for rejecting the null hypothesis with a 5% type I error rate.

### Data Availability

The bulk RNAseq data generated in this study are publicly available at Gene Expression Omnibus (GEO) at [accession number provided prior to publication]. Raw/Processed data are available as Supplemental Data Tables (Suppl. Data Tables 1-8).

---

**Supplementary Figure S1:**
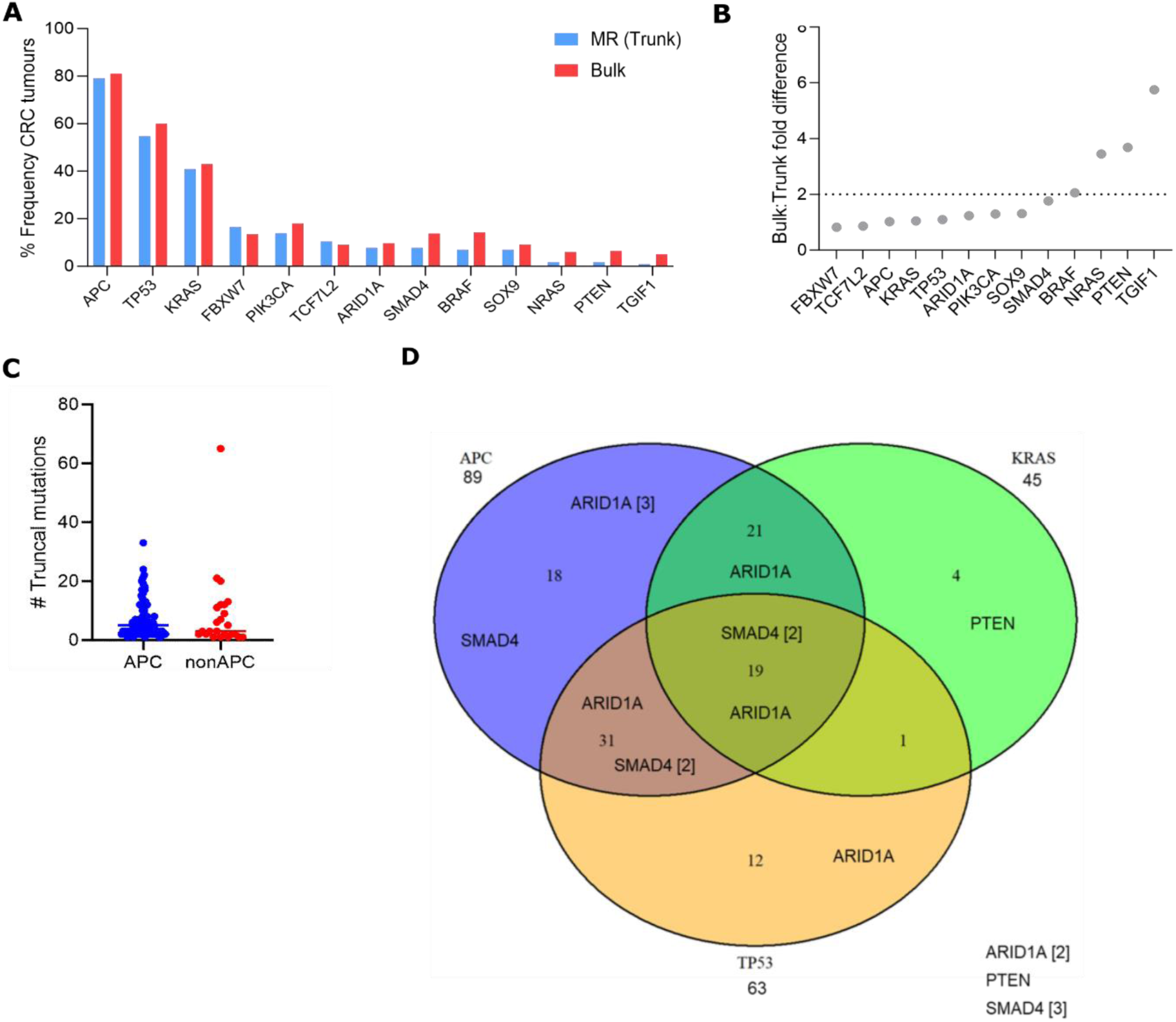
Metanalysis of truncal mutations identified using multi-regional sampling. **(A)** Comparison of mutation frequency from the trunk of phylogenetic trees created from multi-regional studies or bulk sequencing (TCGA data) showing genes with dN/dS >1 (q <0.001) (Martincorena et al., 2017 CRC). **(B)** Fold difference in bulk versus trunk mutation frequency. **(C)** Number of truncal mutations in APC or non-APC driven tumours considering all detected mutations. **(D)** Venn diagram of relationship between APC, TP53 and KRAS driven tumours. PTEN, SMAD4 and ARIDlA mutations annotated. Multiregional studies, N=llS tumours.

**Supplementary Figure S2:**
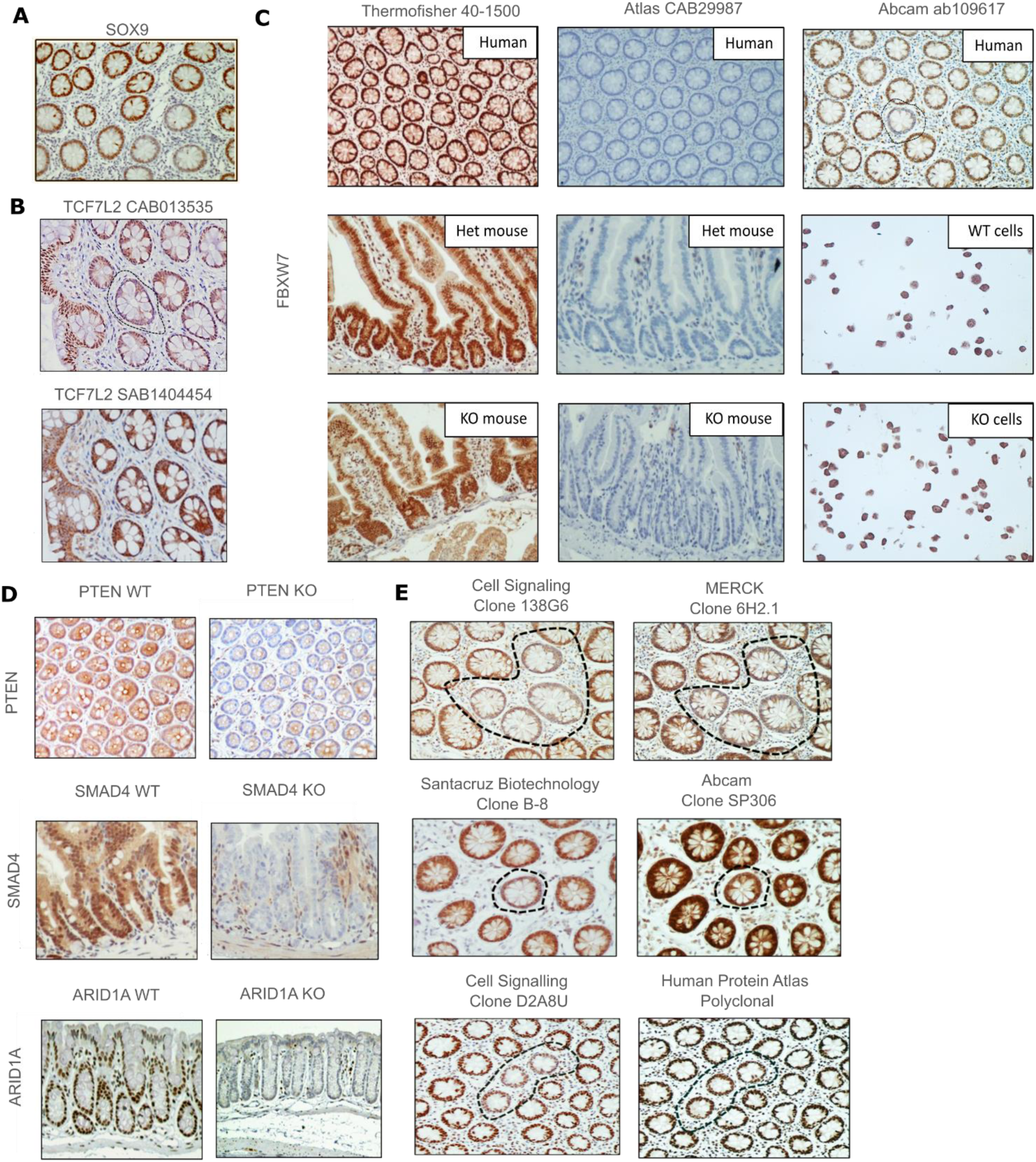
Antibody screen of tumour suppressor genes. **(A)** SOX9 detection failed screening due to non-uniform expression across crypt axis. **(B)** TCF7L2 detection failed screening as identified clone not detected with independent antibody. **(C)** Top row shows staining of FBXW7 antibodies in human tissue. First antibody (Thermofisher) had strong staining but no detecetd clones, second antibody (Atlas CAB) showed no staining and third antibody (Abeam) led to identification of deficient clones. FBXW7 detection failed screening as antibodies did not distinguish known knockout sample in comparison of het and knockout samples. (Mouse: VillinCreERT;FBXW7 mouse model. Human: HEK293T cells with FBXW7 was knocked out using CRISPR/Cas9 and isogenic controls). **(D)** Genetic knockout validation of antibodies used for PTEN. SMAD4 and ARIDlA. VillinCreERT promoter used for to recombine appropriate floxed alleles in mice. **(E)** Independent antibody validation of detected PTEN, SMAD4 and ARIDlA clones.

**Supplementary Figure S3:**
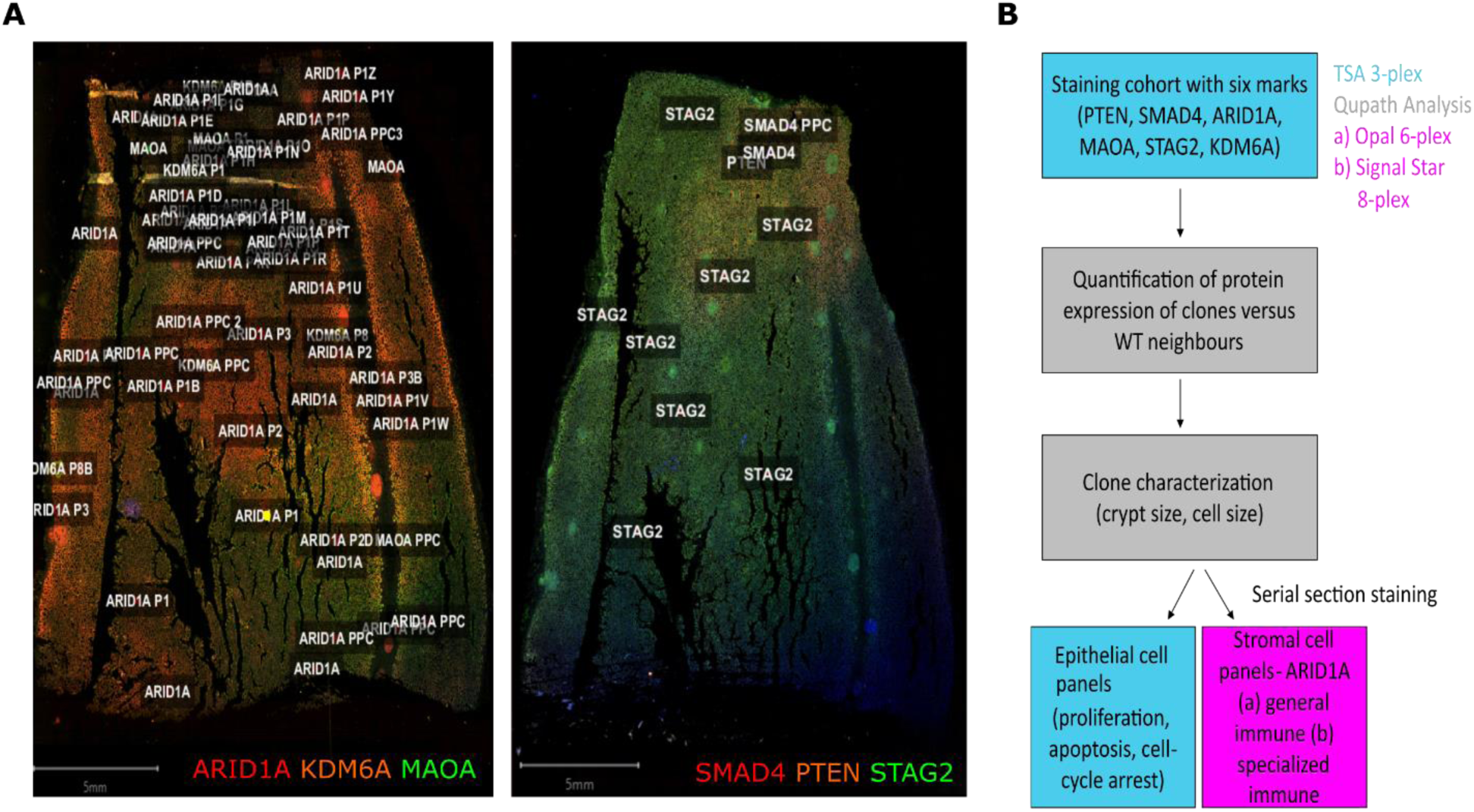
Analysis of tissue sections for multiplexing immunofluorescence. **(A)** Whole slide imaging and annotation of clones. Stained as two 3-plex panels of two serial section shown. **(B)** Schematic of workflow for sections from selected patient cohort and application of multiplexing panels.

**Supplementary Figure S4:**
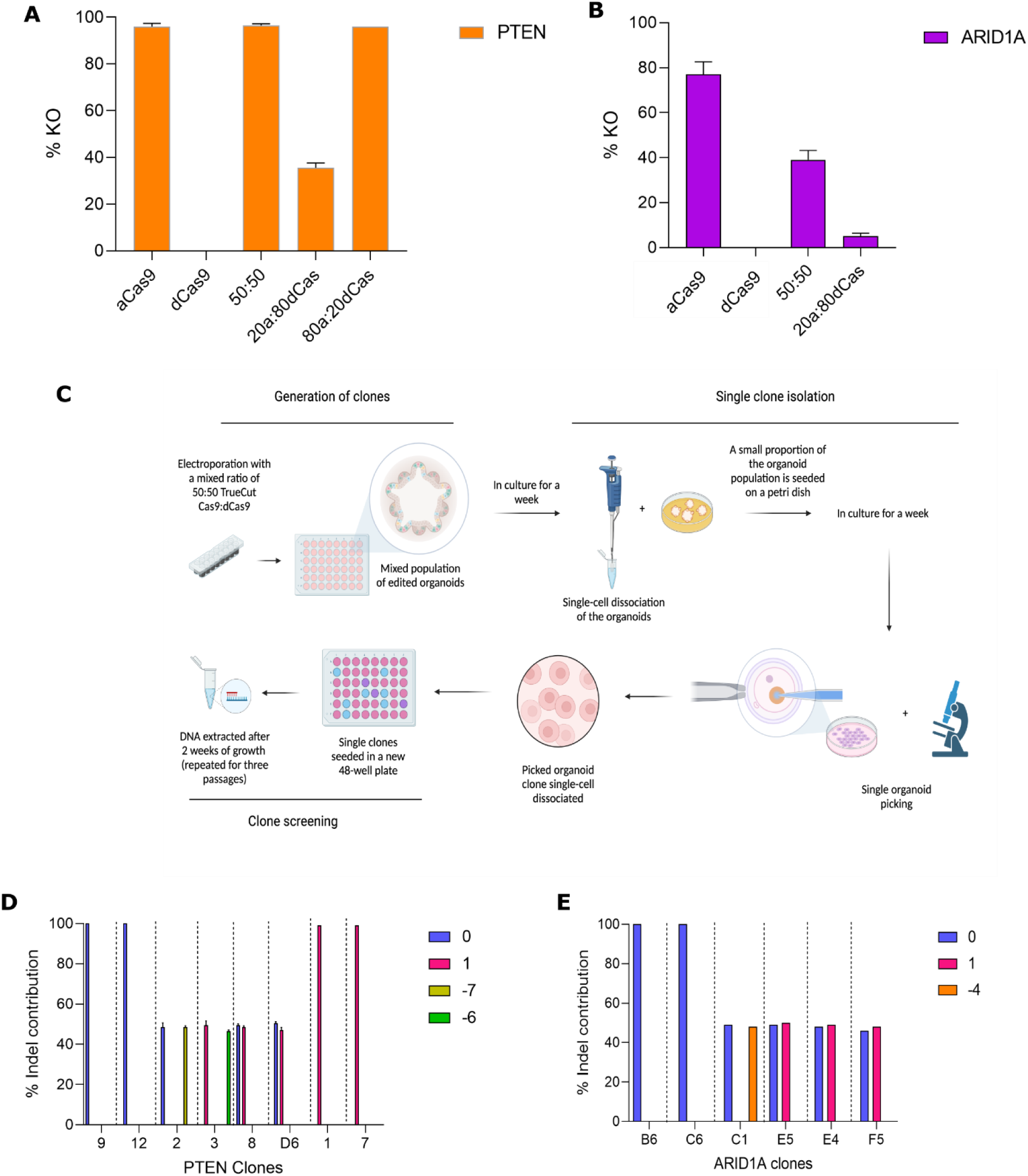
Generation of organoids with heterozygous mutations by CRISPR-Cas9 ribonucleoprotein-based editing. **(A-B)** Optimisation of ratio of active Cas9 (aCas9) to dead Cas9 (dCas9) forPTEN (20a:80dCas) (A) and ARID1A guides (50:50) (B). **(C)** Workflow for generation and isolation of heterozygous organoids. Image created on BioRender. **(D, E)** Editing outcomes showing contribution of alleles for each PTEN (A) and ARIDlA clone (E). 0 indicates WT allele.

**Supplementary Figure S5:**
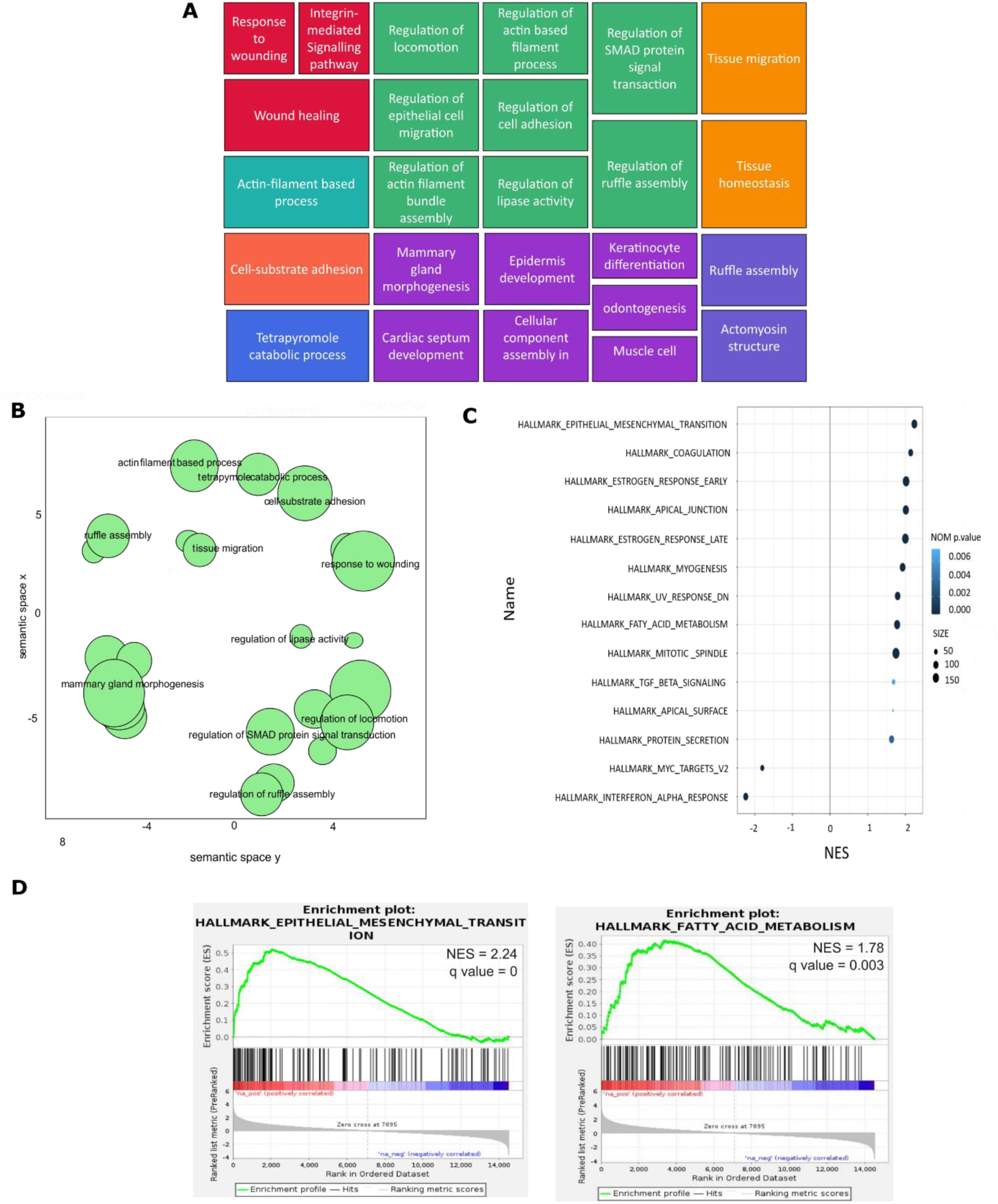
Gene set enrichment analysis {GSEA} for PTEN heterozygous vs WT colonic organoids. **(A)** Gene Ontology Biological Processes (GOBP) coloured based on semantic similarity. Different colours represent different clusters which are interacting between them. Positive enrichment. **(B)** Number of gene sets in clusters grpuped based on semantic similarity. Showing Cluster representatives, which are remaining terms after redudancy removal. Positive enrichment. **(C)** GSEA for hallmark pathways. Considering gene sets FDR q-value < 1%. **{D)** Example of relevant positively enriched hallmark pathways.

**Supplementary Figure S6:**
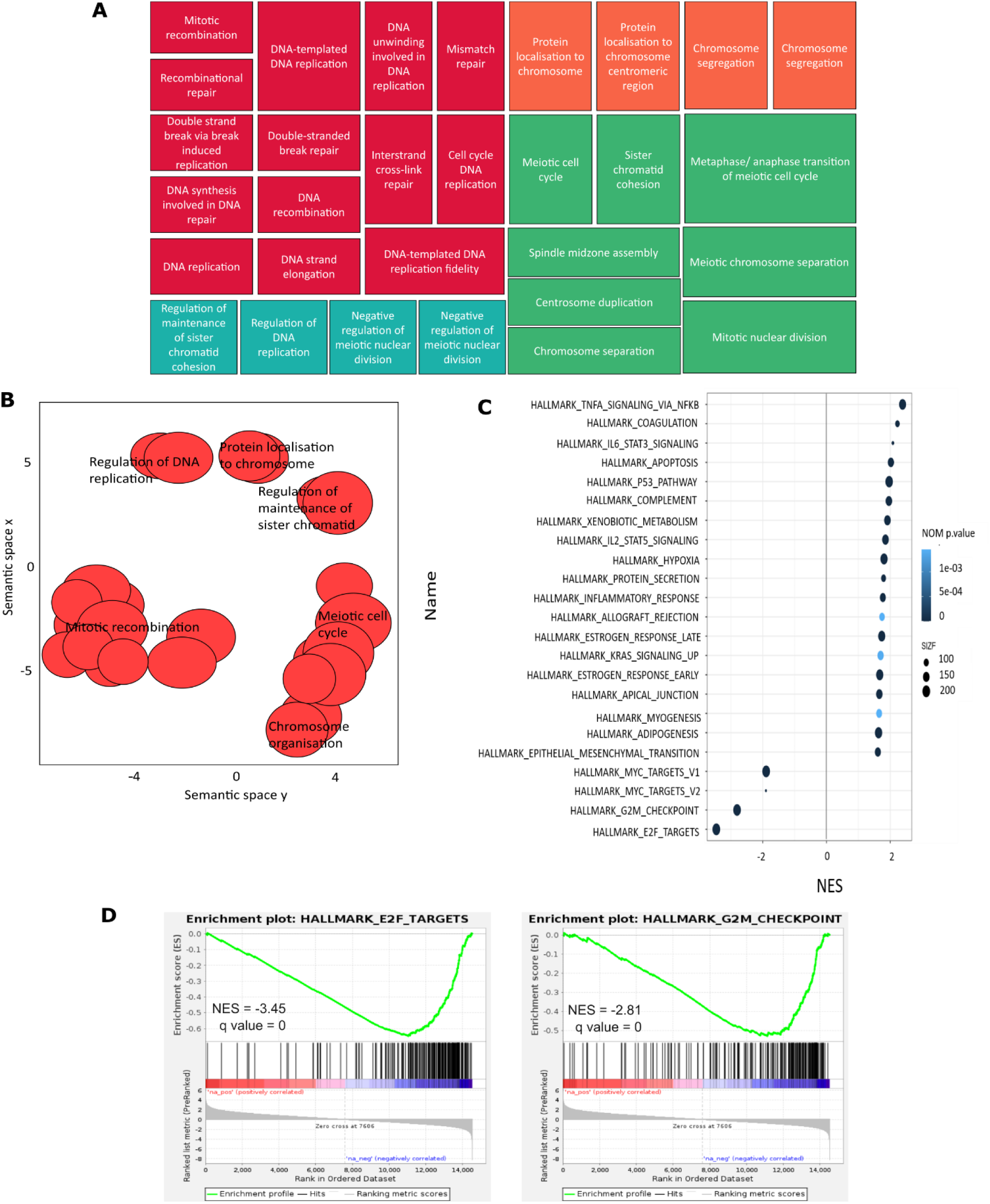
Gene set enrichment analysis (GSEA) for ARID1A heterozygous vs WT colonic organoids. **(A)** Gene Ontology Biological Processes (GOBP) coloured based on semantic similarity. Different colours represent different clusters which are interacting between them. Negative enrichment. **(B)** Number of gene sets in clusters grpuped based on semantic similarity. Showing Cluster representatives, which are remaining terms after redudancy removal. Neagtive enrichment. **(C)** GSEA for hallmark pathways. Considering gene sets FDR q-value < **1%. {D)** Example of relevant positively enriched hallmark pathways.

**Supplementary Figure S7:**
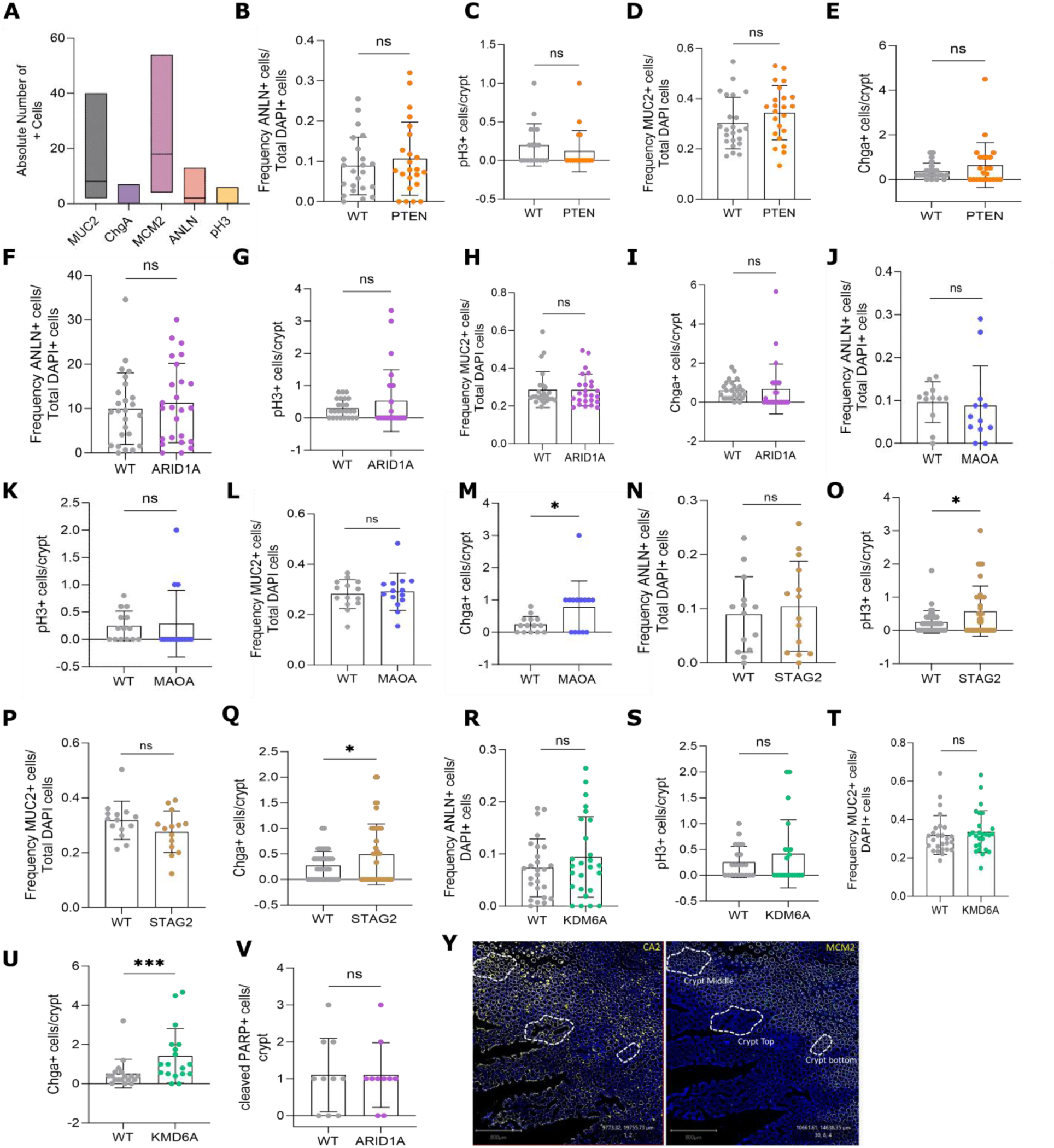
Quantification of proliferation and differentiation markers in deficient clones. **(A)** Number of positive cells per wlldtype crypt for markers shown. **(B-V)** Frequency of cells expressing the markers shown associated with the stated clones. Expressed as frequency of positive cells of total DAPI+ cells (B,D,F,H,J,L,N,P,R,T) or as number of positive cells per crypt (C,E,G,I,K,M,0,Q,S,U,V). **(Y)** Definition of crypt axis levels based on CA2 and MCM2 staining in en face embedded tissue. CA2+ MCM2− crypt top. CA2+ MCM2+ crypt middle. CA2− MCM2+ crypt bottom. Paired t-test or Wllcoxon test was performed to assess statistical significance based on type of distribution. (B) p = 0.297, (C) p = 0.203, (D) p = 0.175, (E) p= 0.355, (F) p = 0.178, (G) p = 0.597, (H) p = 0.56, (I) p = 0.406, (J) p = 0.32, (K) p < 0.999, (L) p = 0.541, (M) p = 0.017, (N) p = 0.36, (O) p = 0.025, (P) p = 0.075, (Q) p = 0.035, (R) p = 0.12, (S) p = 0.508, (T) p = 0.247, (U) p = 0.0004, (V) p < 0.999. * p < 0.05, *** p < 0.001.

**Supplementary Figure S8:**
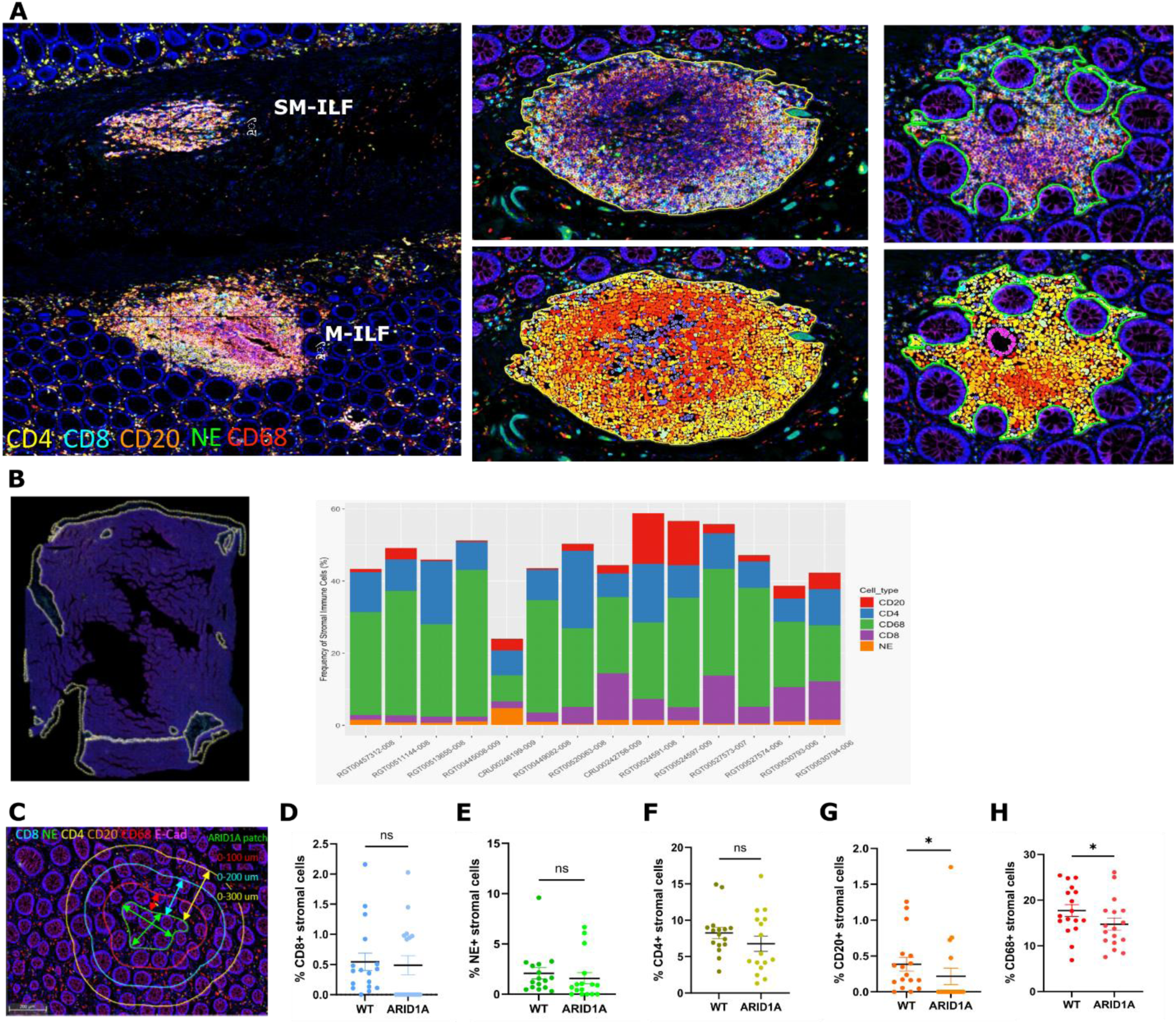
Characterisation of immune cell populations. **(A)** Examples of immune lymphoid follicles in the submucosa (SM-ILF) and the colonic mucosa (M-ILF). Zoomed in images show one example of SM-ILF (left) and one example of M-ILF (right). Bottom pictures show cell segmentation. **(B)** Whole­ section analysis (excluding submucosa) showing contribution of different immune cell types. **(C)** Infiltration analysis within and around ARIDlA patches. **(D-H)** Percentage of positive stromal cells as defined by trained crypt stroma random forest classifier (Halo, Indica labs). Cells measured using cell segmentation algorithm within ARID1A patch (>5 crypts) or 300 um radius around the patch (WT). CD8+ cytotoxic T-cells (D). NE+ neutrophils (E). CD4+ T-helper cells(F). CD20+ B-cells (G). CD68+ macrophages (H). Paired Wilcoxon test was performed to assess statistical significance. (D) p = 0.528, (E) p = 0.175, (F) p = 0.065, (G) p = 0.030, (H) p = 0.027. * p < 0.05.

**Supplementary Figure S9:**
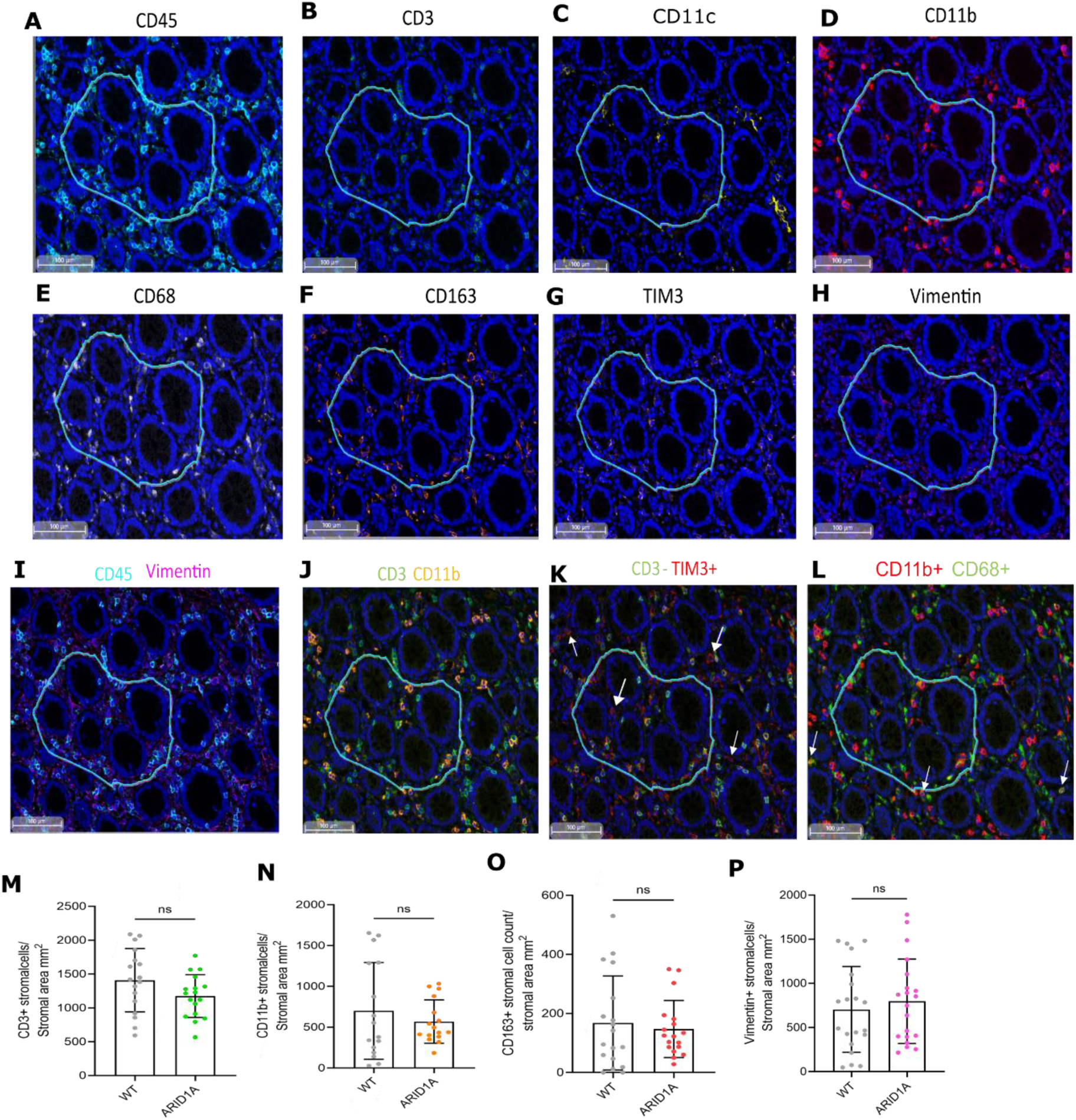
Signal star stromal cell profiling of cell populations within and around ARID1A patches. **(A-H)** Individual markers around an ARIDlA patch shown in blue. Arranged in order of staining. **(I)** Separation of immune cells (CD45+) and mesenchymal cells (Vimentin+). **(J)** Separation of lymphocytic T-cells (CD3+) and myeloid cells (CDllb+). **(K)** TIM3+/CD3-cells indicated by the arrows. **(L)** CD11b+/CD68+ cells indicated by the arrows. **(M-P)** Quantification of stromal cells in WT and AR!DlA deficient patches. Showing non-significantly different stromal populations. CD3+ cells (M). CDllb+ cells (N). CD163+ cells (0). Vimentin+ cells (P). Paired Wilcoxon test was performed to assess statistical significance. **(M)** p =0.0638, (N) p = 0.495, (0) p = 0.766, (P) p = 0.648.

**Table S1:**
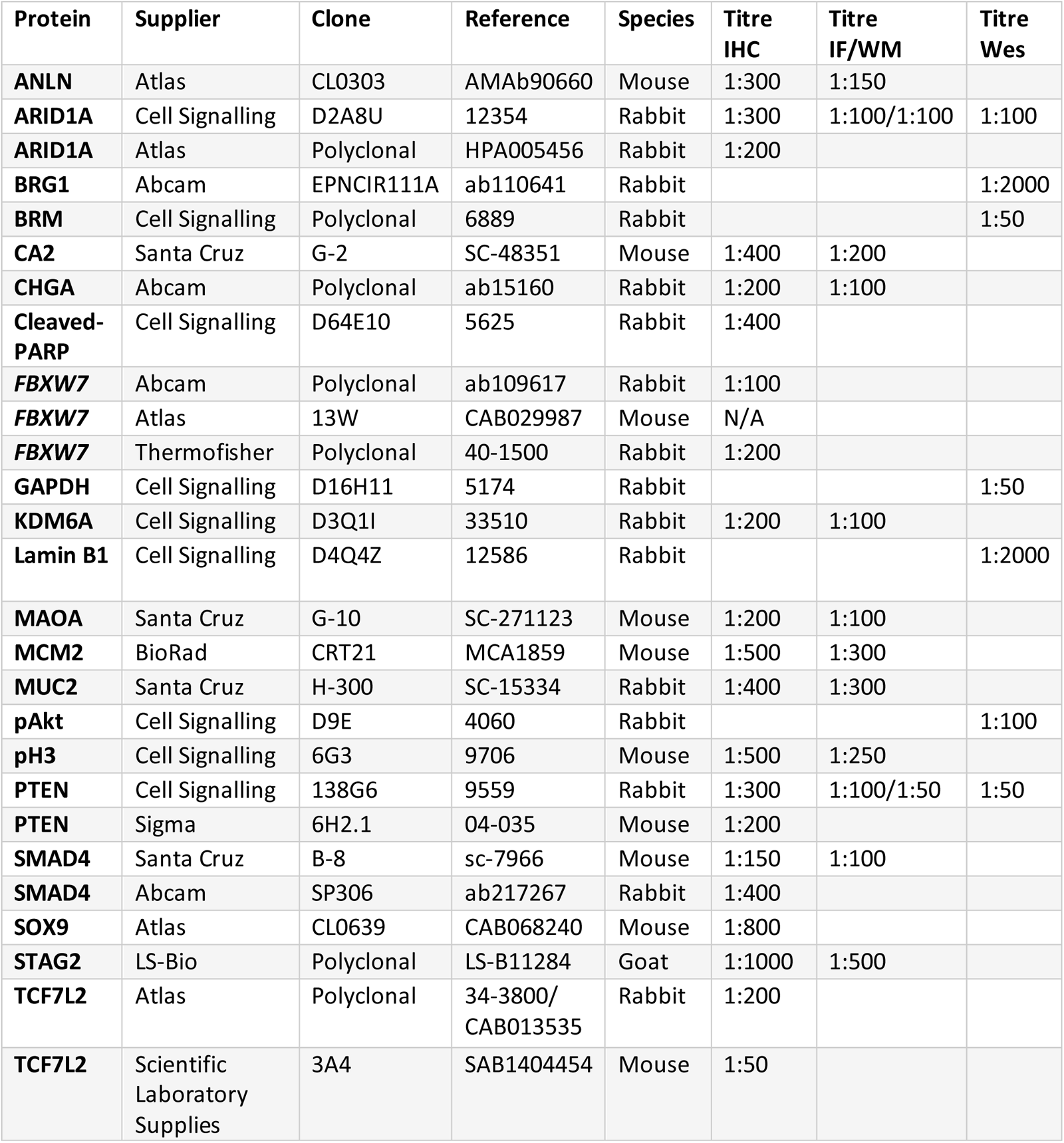
Primary antibodies used for staining. *N/A indicates that no suitable titre was found in human and mouse colonic tissue. IHC: immunohistochemistry, IF: immunofluorescence, WM: Whole-mount*.

| Protein | Supplier | Clone | Reference | Species | Titre IHC | Titre IF/WM | Titre Wes |
| --- | --- | --- | --- | --- | --- | --- | --- |
| **ANLN** | Atlas | CL0303 | AMAb90660 | Mouse | 1:300 | 1:150 |  |
| **ARID1A** | Cell Signalling | D2A8U | 12354 | Rabbit | 1:300 | 1:100/1:100 | 1:100 |
| **ARID1A** | Atlas | Polyclonal | HPA005456 | Rabbit | 1:200 |  |  |
| **BRG1** | Abcam | EPNCIR111A | ab110641 | Rabbit |  |  | 1:2000 |
| **BRM** | Cell Signalling | Polyclonal | 6889 | Rabbit |  |  | 1:50 |
| **CA2** | Santa Cruz | G-2 | SC-48351 | Mouse | 1:400 | 1:200 |  |
| **CHGA** | Abcam | Polyclonal | ab15160 | Rabbit | 1:200 | 1:100 |  |
| **Cleaved-PARP** | Cell Signalling | D64E10 | 5625 | Rabbit | 1:400 |  |  |
| **FBXW7** | Abcam | Polyclonal | ab109617 | Rabbit | 1:100 |  |  |
| **FBXW7** | Atlas | 13W | CAB029987 | Mouse | N/A |  |  |
| **FBXW7** | Thermofisher | Polyclonal | 40-1500 | Rabbit | 1:200 |  |  |
| **GAPDH** | Cell Signalling | D16H11 | 5174 | Rabbit |  |  | 1:50 |
| **KDM6A** | Cell Signalling | D3Q1I | 33510 | Rabbit | 1:200 | 1:100 |  |
| **Lamin B1** | Cell Signalling | D4Q4Z | 12586 | Rabbit |  |  | 1:2000 |
| **MAOA** | Santa Cruz | G-10 | SC-271123 | Mouse | 1:200 | 1:100 |  |
| **MCM2** | BioRad | CRT21 | MCA1859 | Mouse | 1:500 | 1:300 |  |
| **MUC2** | Santa Cruz | H-300 | SC-15334 | Rabbit | 1:400 | 1:300 |  |
| **pAkt** | Cell Signalling | D9E | 4060 | Rabbit |  |  | 1:100 |
| **pH3** | Cell Signalling | 6G3 | 9706 | Mouse | 1:500 | 1:250 |  |
| **PTEN** | Cell Signalling | 138G6 | 9559 | Rabbit | 1:300 | 1:100/1:50 | 1:50 |
| **PTEN** | Sigma | 6H2.1 | 04-035 | Mouse | 1:200 |  |  |
| **SMAD4** | Santa Cruz | B-8 | sc-7966 | Mouse | 1:150 | 1:100 |  |
| **SMAD4** | Abcam | SP306 | ab217267 | Rabbit | 1:400 |  |  |
| **SOX9** | Atlas | CL0639 | CAB068240 | Mouse | 1:800 |  |  |
| **STAG2** | LS-Bio | Polyclonal | LS-B11284 | Goat | 1:1000 | 1:500 |  |
| **TCF7L2** | Atlas | Polyclonal | 34-3800/<br>CAB013535 | Rabbit | 1:200 |  |  |
| **TCF7L2** | Scientific Laboratory Supplies | 3A4 | SAB1404454 | Mouse | 1:50 |  |  |

**Table S2:** Multiplex IF panels. Clonal fates cell cycle panel was designed with two stripping cycles where at the second cycle two antibodies of different species were added, CA2 was added with Tyramide Signal Amplification reagent and pH3 with Alexa Fluorophore.

| Panel name | Targets order | Fluorophore order | Multiplexing method |
| --- | --- | --- | --- |
| Clonal Marks-1 | SMAD4 PTEN STAG2 | Cy5, Cy3, 488 | Standard TSA |
| Clonal Marks-2 | ARID1A KDM6A MAOA | Cy5, Cy3, 488 | Standard TSA |
| Epithelial clonal properties lineage | MUC2 MCM2 ChgA | Cy5, Cy3, 488 | Standard TSA |
| Epithelial clonal properties cell cycle | ANLN CA2+pH3 | Cy5, Cy3 + 488 | Standard TSA |
| General Immune cells | CD8 NE CD4 CD20 CD68<br>Ecadherin | 480, 520, 570, 620, 690, 780 | Opal TSA |
| Specialized Immune cells | Round 1: CD45 CD3 CD11c<br>CD11b<br>Round 2: CD68 CD168 TIM3<br>Vimentin | 488, 594, 647, 750 | Signal Star |

**Table S3:**
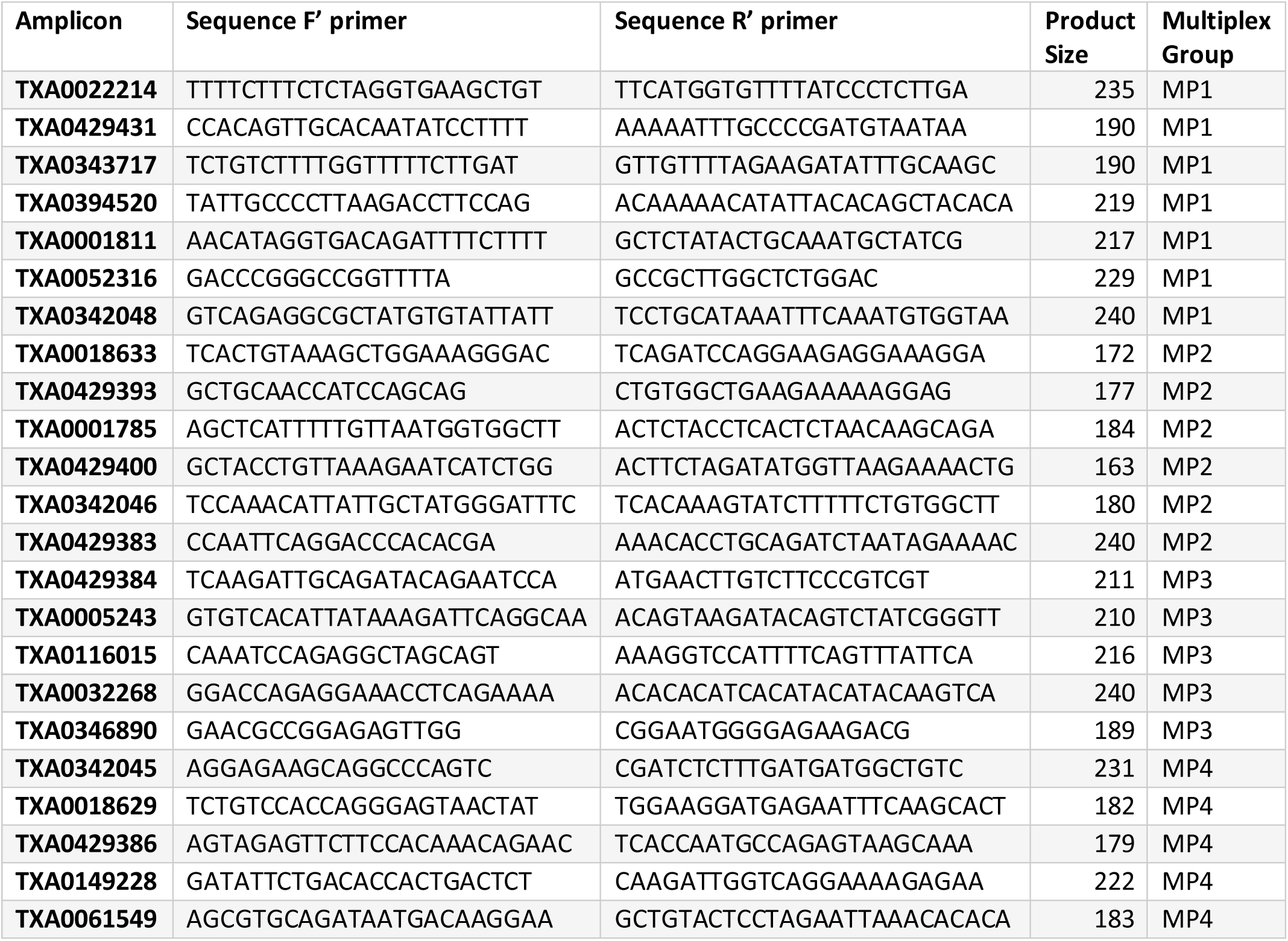
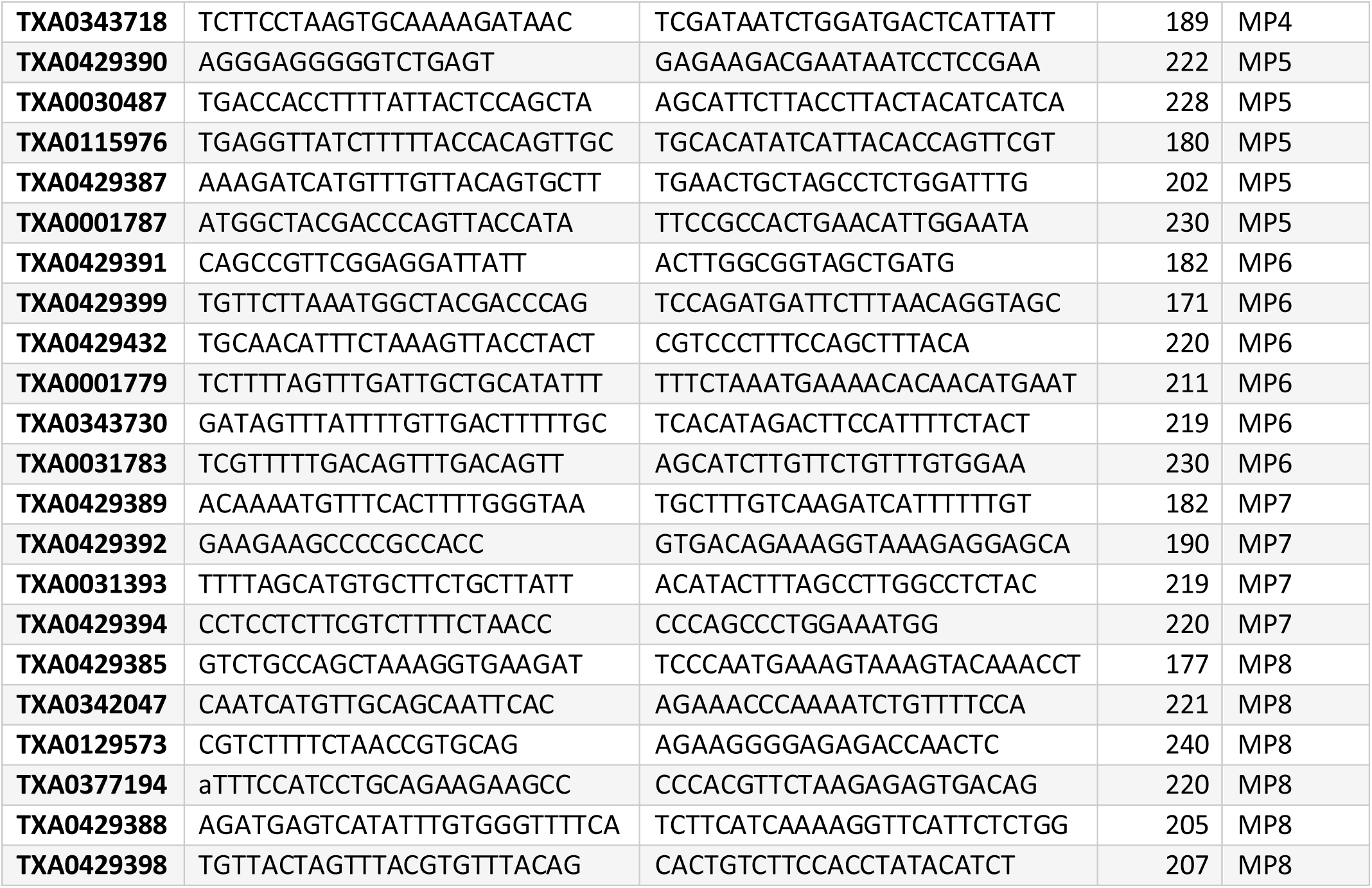
Fluidigm Primers for PTEN exon coverage used for Juno chip and pooling strategy. Lower case letters indicate intron binding.

**Table S4:**
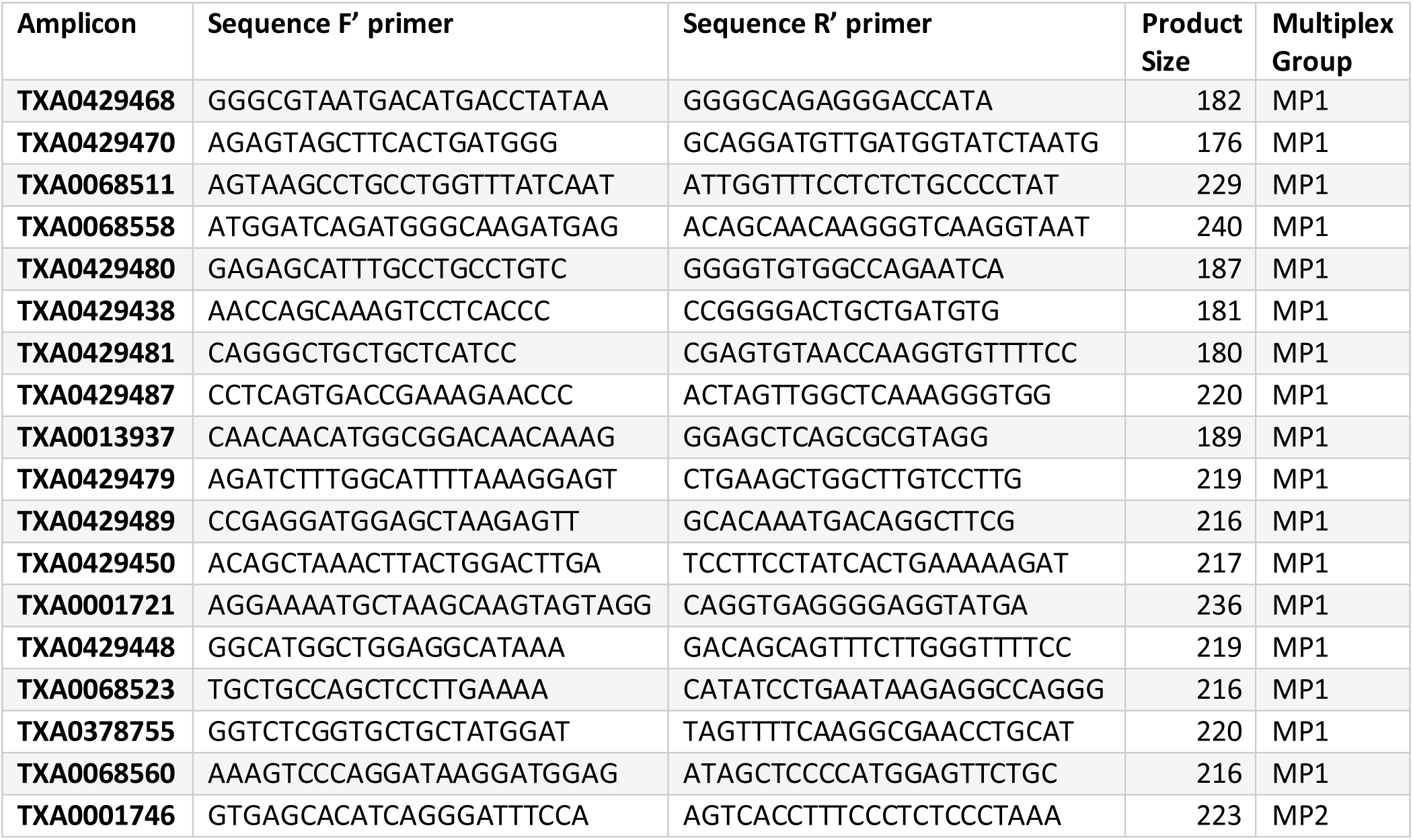
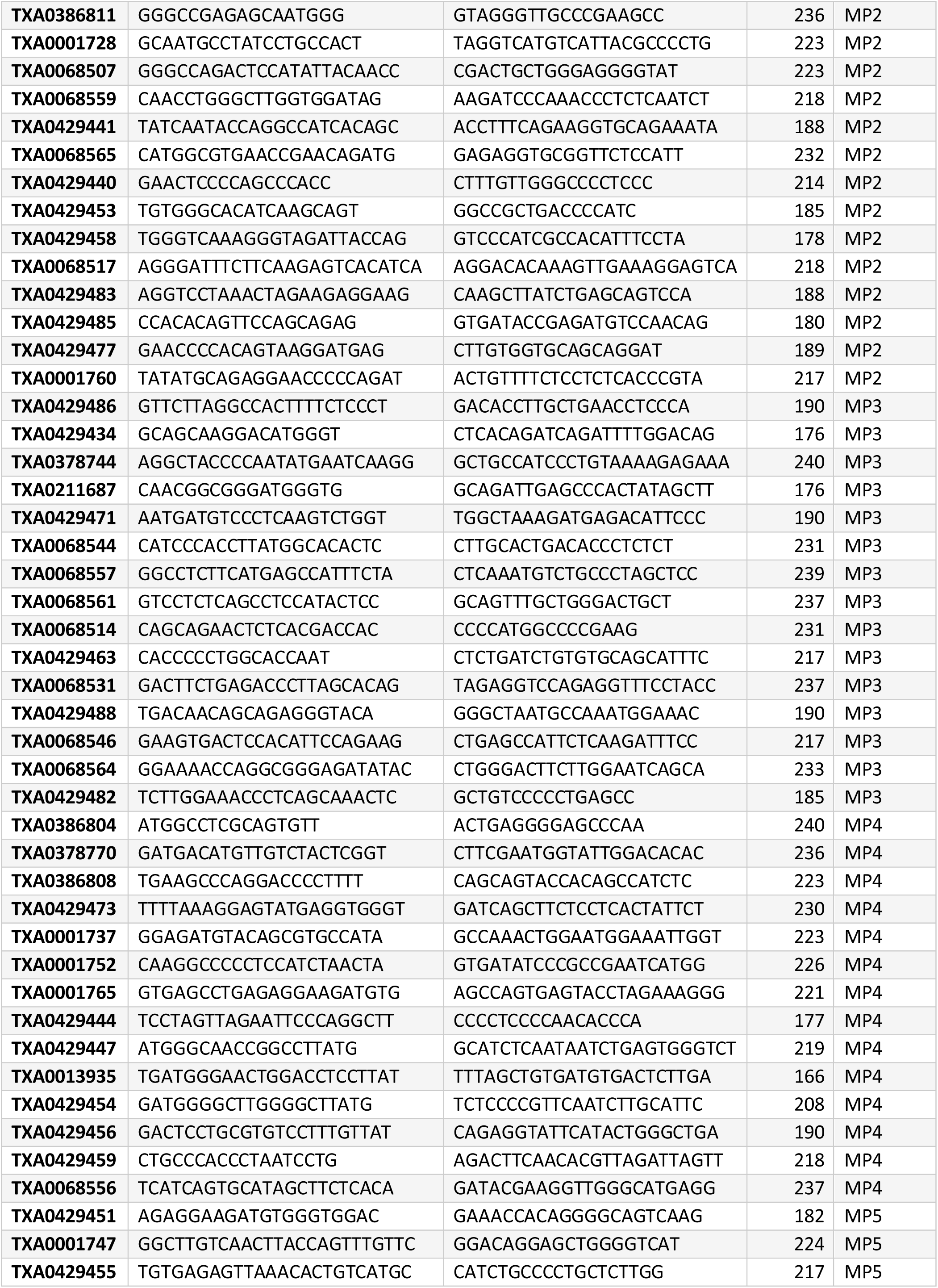
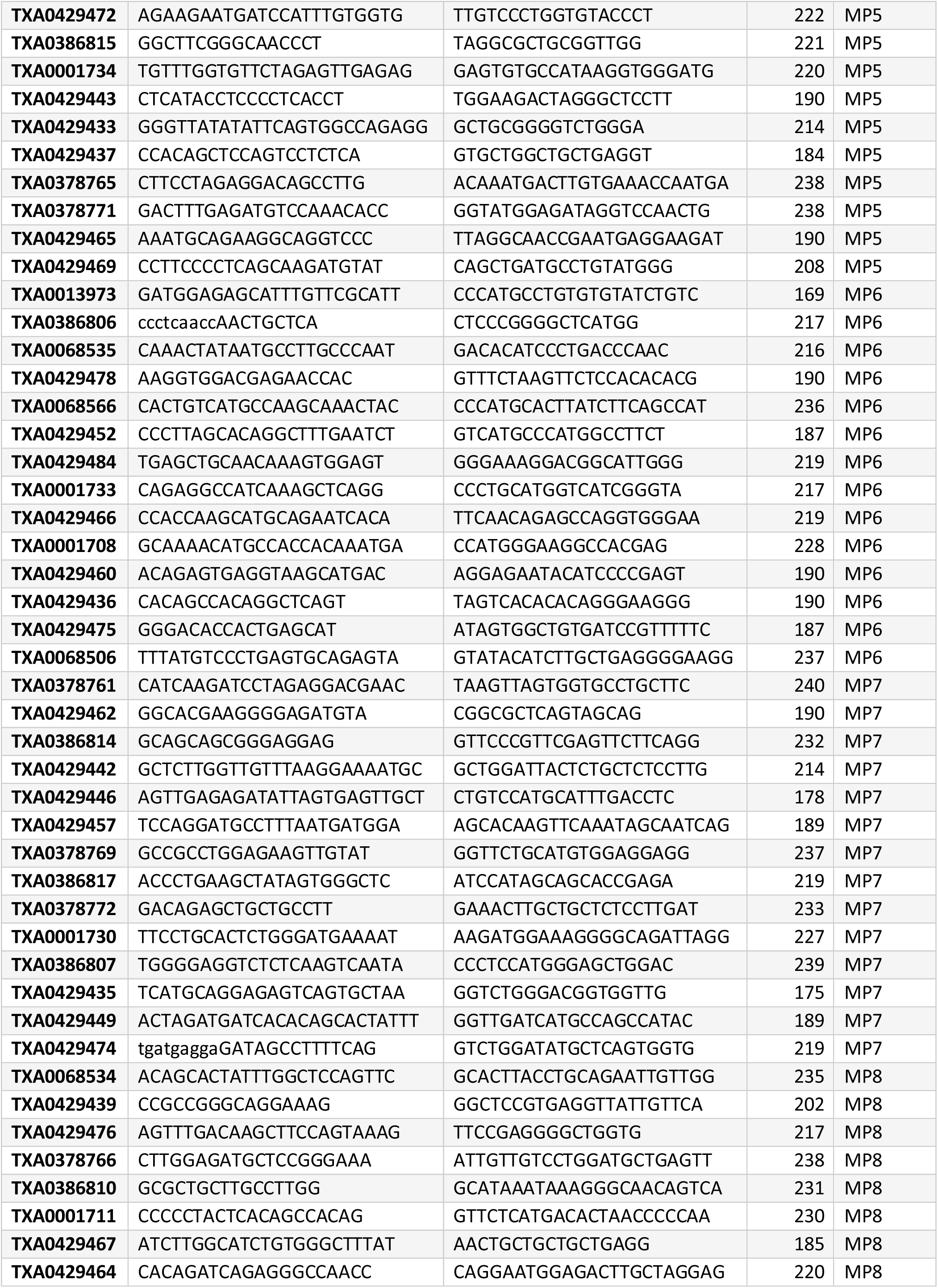
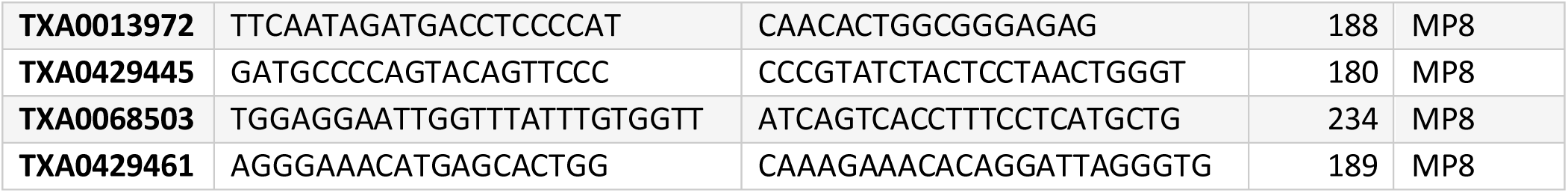
Fluidigm Primers for ARID1A exon coverage used for Juno chip and pooling strategy. Lower case letters indicate intron binding.

